# Mobile EEG for Neurourbanism Research - What Could Possibly Go Wrong? A Critical Review with Guidelines

**DOI:** 10.1101/2024.03.22.586309

**Authors:** Klaus Gramann

## Abstract

Based on increasing incidents of mental ill-health associated with living in dense urban environments, the field of Neurourbanism developed rapidly, aiming at identifying and improving urban factors that impact the health of city dwellers. Neurourbanism and the closely related field of Neuro-Architecture have seen a surge in studies using mobile electroencephalography (EEG) to investigate the impact of the built and natural environment on human brain activity moving from the laboratory into the real world. This trend predominantly arises from the ready availability of affordable and portable consumer hardware, which not only guarantees operational simplicity but also frequently incorporates automated data analysis functions. This significantly streamlines the process of EEG data acquisition, analysis, and interpretation, seemingly challenging the necessity of specialized expertise in the method of EEG or neurosciences in general. As a consequence, numerous studies in the field of Neurourbanism have used such off-the-shelf systems in laboratory and real-world experimental protocols including active movement of participants through the environment. However, the recording and analysis of EEG data entails numerous requisites, the disregard of which may culminate in errors during data acquisition, processing, and subsequent interpretation, potentially compromising the scientific validity of the outcomes. The often relatively low number of electrodes offered by affordable and portable consumer EEG systems further restricts specific analyses approaches to the low-dimensional EEG data. Crucially, a large part of Neurourbanism studies used black-box analyses provided by such consumer systems or incorrectly applied complex data-driven analyses methods that are incompatible with the recorded low-dimensional data. The current manuscript delineates the prerequisites concerning EEG hardware and analytical methodologies applicable to stationary and mobile EEG protocols, whether conducted within a controlled laboratory environment or in real-world settings. It conducts a comprehensive review of EEG studies within the domain of Neurourbanism and Neuro-Architecture, assessing their adherence to these prerequisites. The findings reveal severe deficiencies in the utilization of hardware and data processing methods, thereby rendering these studies unsuitable for scientific scrutiny. Consequently, the present paper provides guidelines for the selection of EEG hardware and analytical strategies for researchers engaged in mobile EEG recordings, be it within a laboratory or real-world context, aimed at steering future investigations in the field of Neurourbanism and Neuro-Architecture.

## Background

Over the last century, a rapidly growing urbanization took place leading a large part of the world’s population to move to and take advantage of the resources of urban centers. This trend, however, was accompanied by increased risks of physical and mental ill-health for city dwellers (Kennedy & Adolphs, 2011). The adverse effects of urbanization have spurred the emergence of Neurourbanism as a research discipline that concentrates on the welfare of urban residents by identifying and investigating the factors of urban living that influence their health (Adli et al., 2017). Neurourbanism proposes an interdisciplinary approach using a wide variety of methods to gain a better understanding of the human response to the built environment. The promise of “neuro” in Neurourbanism is to incorporate neuroscientific measures besides the existing assessment tools to provide more “objective” insights into the subjective responses to specific aspects of the urban environment and how these relate to well-being. With the initial work of Ulrich (Ulrich, 1981) and Eberhard (Eberhard, 2009a, 2009b), the use of neuroscientific methods to specifically investigate the brain responses of humans to the built environment has seen a constant increase with electroencephalography (EEG), functional magnetic resonance imaging (fMRI), and functional near-infrared spectroscopy (fNIRS) as the leading methods in this field (Ancora et al., 2022).

### Neurourbanism from the Lab to the Real World

Studying the brain response to the built environment in situ, however, is limited by the restrictions inherent in many established neuroimaging methods mainly due to the weight of the systems and their susceptibility to movement-artifacts (Makeig et al., 2009). Further, different brain imaging methods assess different aspects of brain activity. fMRI and fNIRS measure hemodynamic activity (i.e., changes in blood flow in different brain regions) with high spatial resolution but relatively slow temporal resolution due to the sluggish nature of blood flow. In contrast, EEG measures the electrical activity of cortical neurons with high temporal resolution but relatively low spatial resolution due to the unknown mixture of volume- and capacitive conducted signals that mix at the sensor level (Gramann, Jung, et al., 2014; Mehta & Parasuraman, 2013). The use of MRI is expensive and access often limited, the systems are heavy, and the recorded signal is susceptible to movements requiring stationary protocols with participants lying supine in the scanner not allowed to move at all (Gramann et al., 2011). Experimental protocols using MRI are thus restricted to watching images or videos of the built environment while the associated brain dynamics are recorded (e.g., Kim et al., 2010; Kühn et al., 2021). It is possible though to combine information about the individual living conditions and responses to urban or other stressors beyond the immediate response to urban stimulus material (Dimitrov-Discher et al., 2023; Spiers & Maguire, 2007). EEG is relatively inexpensive, the systems are smaller and often portable but the recordings are also susceptible to motion artifacts. Thus, Neurourbanism studies using EEG as a method to assess neural responses to the built or natural environment often use stationary protocols (e.g., Grassini et al., 2019; Mahamane et al., 2020). These traditional EEG protocols reduce behavioral responses of participants to a bare minimum - often button presses at the end of a trial - to avoid movement-related artifacts from contaminating the feeble signals of interest (Gramann, Ferris, et al., 2014). Such highly controlled laboratory-based studies provide the advantage of control over all factors of interest, which especially for research in urban contexts might otherwise not be controllable, including traffic, ambient noise, people, or weather conditions (Vallet & Van Wassenhove, 2023).

However, since human beings naturally interact with their built and social surroundings and experience the world by moving through and interacting with it, traditional stationary human brain imaging protocols are often criticized as being artificial due to their sensorial, social, and contextual deprivation (Shamay-Tsoory & Mendelsohn, 2019). It seems further questionable whether watching 2D representations of the real world while sitting in a sparsely illuminated lab or lying in a scanner share comparable affordance with the built environment in situ (Djebbara & Kalantari, 2023). As a result, criticisms frequently arise against laboratory protocols for yielding ecologically invalid results, particularly when the targeted cognitive or behavioral processes lack clear definitions (Holleman et al., 2020), and this might be especially the case for the field of Neurourbanism.

To enhance ecological validity and address the limitations associated with stationary laboratory protocols, researchers have increasingly explored human brain activity in relation to the built or natural environment in actively behaving participants in the laboratory or in real-world settings. Over the past decade, there has been a notable rise in publications within the field of Neurourbanism (Ancora et al., 2022), accompanied by a surge in real-world mobile EEG investigations in general (Niso et al., 2023). The emergence of low-cost, compact, and portable EEG systems seemingly capable of recording neural activity during active movement inside or outside traditional laboratories has opened new avenues for studying human neural responses to the built or natural environment, offering the potential to overcome the ecological validity constraints associated with conventional laboratory research. The affordable cost, ready availability off-the-shelf, ease of application, and automated data analysis that is sold with these systems suggest that researchers from various disciplines can utilize EEG to gain deeper insights into the human response to the environment even without neuroscience training or specific EEG expertise. As a consequence, the number of Neurourbanism studies using EEG and mobile EEG in the laboratory or the real-world increased significantly over the last decade.

### The Pitfalls of (mobile) EEG

While the benefit of mobile neuroscientific methods for Neurourbanism research, specifically mobile brain imaging methods seems obvious, the necessary expertise for recording scientifically valid data in the lab or the real world, as well as the subsequent analyses and interpretation of complex and high-dimensional data, is often less emphasized. Leveraging commercially available EEG systems with modest costs, assuring easy setup and operational simplicity, along with algorithms facilitating automated analyses of EEG data and classification of human states, appears to present a facile solution to mitigate the need for specialized expertise. Nevertheless, publications in the field of Neurourbanism based on such systems often seem to encounter challenges in conducting methodologically sound and replicable neuroscientific studies. Incorrect application of the methods and assumptions of what the system can and cannot do in combination with inadequate or even black box data analysis approaches can lead to drastic misinterpretation of the outcome. With an increasing number of predatory journals that favor profit over scientific quality, such results will still be published and can have lasting effects on the field.

The present review thus addresses critical issues in the new and fast-growing field of Neurourbanism studies that use EEG or mobile EEG protocols providing an overview of the method and analyses. In the end, guidelines will be provided for EEG protocols in general and specifically for mobile EEG protocols in Neurourbanism research as these require additional considerations regarding hardware and analyses approaches. In the subsequent sections, the primary challenges associated with employing EEG recordings in the laboratory or real-world experiments are delineated. Commencing with a brief overview of the origin of the EEG signal, an examination of crucial considerations regarding the recording hardware and the analytical approaches follows. The focus will be on two main aspects that are critical for EEG studies in general and specifically for real-world neuroscientific studies using mobile EEG: i) The recording equipment, encompassing amplifier specifications, and details on the quantity, type, and application of electrodes, and ii) EEG analyses, including data preprocessing and downstream extraction of brain electrical features. These aspects are evaluated based on publication guidelines and recommendations for EEG studies (Keil et al., 2014). After describing the methodological foundations regarding the EEG recording technology and analysis approaches, publications in the domain of Neurourbanism using EEG will be analyzed with respect to the described methodological issues with a special focus on EEG studies conducted in the real world or using mobile EEG methods. Summarizing the methodological requirements and the results from the literature review, guidelines will be provided allowing to narrow down equipment options and analytical protocols for Neurourbanism studies with more ecologically valid protocols.

### EEG Origins, Volume Conduction, and Capacitive Conduction

The EEG signal mainly originates from the synchronized electrical activity of thousands to millions of pyramidal neurons located in the human cortex with a perpendicular orientation of these neurons to the surface of the cortex (Lopes da Silva, 2013). The EEG signal is based on postsynaptic activity in larger populations of these pyramidal neurons resulting in the summed activity of excitatory (EPSPs) or inhibitory (IPSPs) postsynaptic potentials. The specific architecture and the synchronous activation of these neurons produce potentials that are strong enough to be sensed outside the brain volume. The movement of charged particles throughout the brain volume is known as volume conduction. Beyond the brain volume, the potentials extend through capacitive conductance across the protective layers of the meninges (the protective membranes surrounding the brain), skull, and skin to reach the electrodes (Jackson & Bolger, 2014; Nunez & Srinivasan, 2006). At the electrode level, the signals are then conducted through electrolyte to the conductive layer of the electrode, where they are detected and transmitted to the amplifier for amplification of the miniscule signal.

The recorded EEG data typically consist not only of brain activity. Brain signals are characterized by their small amplitudes, typically measured in microvolts (one-millionth of a volt), that have to pass through the less conductive layers of the human meninges, skull, and skin, undergoing spatial dispersion and filtering of high-frequency components. Additional biosignals such as eye and muscle movements, which produce signals with much larger amplitudes, will mix with the signals originating in the brain. These physiological non-brain signals do not encounter the same degree of distortion as brain signals because they don’t traverse the less conductive layers of the meninges and skull. In addition to physiological non-brain signals which can be informative with respect to participants behavioral and psychological state, electronic and mechanical artifacts contribute to the recorded signal. Electronic devices are ubiquitous in the environment introducing interference into the recordings that might lack both spatial stability with respect to the sensors and consistency in their electromagnetic spectrum. Finally, the weight of the amplifier and electrode cables can lead to movement of the system or parts of the system that introduces mechanical artifacts to the recording.

Ecologically more valid protocols allowing for active behaviors in the real world will likely include all of the above described physiological and non-physiological contributions. Active movements of the eyes and muscles, potential system and cap displacements due to head movements of the participants, cable sway during walking as well as external electronic devices in the environment can lead to a different signal being recorded compared to traditional EEG protocols in the laboratory. It is thus imperative that the activity from non-brain sources including biologically relevant activity like eye movement and muscle contraction as well as mechanical and electrical artifacts are dissociated from the brain signal of interest during data analysis. This can be achieved, to a reasonable degree, by good-quality EEG equipment and adequate analytical approaches.

### EEG Recording Equipment

#### Amplifiers Specifications

Over the last decade, a wide range of mobile and wireless EEG amplifiers became available on the market (for an excellent overview of some recent systems see Niso et al., 2023). For Neurourbanism researchers focused on capturing EEG data in the real world, crucial amplifier aspects include the amplifier resolution in bits, the sampling rate in Hertz, and effective common mode rejection (CMR). Additionally, in mobile EEG systems, factors such as the system weight, wireless protocols, and integrated motion sensors are important.

Most commercially available systems offer amplifiers with a resolution ranging from 12 to 24-bit, enabling digitization of a broad signal range of the recorded analog brain activity. Higher resolutions are preferable since they can resolve the signal range in more detail. Similarly for the temporal representation of the signal, higher sampling rates allowing better temporal resolution of the analog signal. As per the Nyquist-Shannon sampling theorem, the sample rate sets the limit for the highest representable frequency in a digitized signal, which is half the sampling frequency. If the frequencies of interest in the recorded EEG signal for a specific application are below 64 Hz, a sampling frequency of 128 Hz would prove sufficient and most commercially available systems typically enable digitization of the analog signal at 128 Hz or higher. The effective CMR, in contrast, varies drastically between available systems ranging from 75 dB to 140 dB. The CMR in an EEG amplifier refers to the amplifier’s ability to minimize signals that are present in both the active and reference electrodes, which are typically noise or interference. CMR thus helps to eliminate unwanted background signals and ensures that the EEG amplifier primarily amplifies the brain’s electrical activity while rejecting common noise sources. For EEG applications, a CMR of 80 dB or higher is often considered good, but the specific requirements can depend on the nature of the recording environment. Because real-world environments come with uncontrollable and often changing ambient electrical noise, higher CMR values are desirable for a mobile EEG amplifier. In terms of weight, the majority of available mobile EEG systems are compact and lightweight, enabling extended data recording sessions without causing participant fatigue. These systems typically employ various industrial wireless protocols for transferring data to mobile devices. Additionally, some systems provide the option to store recorded data locally, necessitating subsequent offline synchronization with other potential data sources. Furthermore, the incorporation of motion sensors into the amplifier systems offers researchers the ability to utilize motion-related information for signal quality control and, in some cases, to identify and remove artifacts associated with movement (e.g., Gwin et al., 2010).

#### Quantity, Types, and Placement of Electrodes

Frequently overlooked but equally significant as amplifier specifications are the considerations surrounding the quantity and types of electrodes employed in mobile EEG recordings. The number of electrodes can vary substantially among different systems. In cases where the primary objective is to reliably extract specific EEG features that are known a priori, the suitable count and positioning of electrodes are determined by the minimum requirements for such extraction. However, in the realm of basic research, particularly within emerging disciplines that often explore novel phenomena previously uninvestigated like in the case of Neurourbanism, our understanding of how the human brain responds to experimental manipulations may be limited. This limitation extends to replicating EEG features established under controlled lab conditions, which might undergo alterations due to participants’ movements during mobile EEG recordings. Consequently, in nascent research domains, the parameters of interest, their spatial distribution across the scalp, and their temporal dynamics are largely unknown or have not yet been adequately replicated to attain validation as reliable parameters. In such circumstances a higher density of the montage with a greater number of electrodes uniformly distributed across the entire scalp becomes indispensable. This approach facilitates the extraction of potential EEG features that co-vary with the experimental manipulation. Importantly, it further permits various analytical methods to distinguish brain activity from non-brain activity. Thus, more electrodes with a uniform distribution across the scalp are better even though they require longer preparation times and are more cumbersome for participants to carry.

Besides the density of the electrode montage, the *kind of electrodes* used can have a major impact on the recorded signal quality. In Neurourbanism research, commonly employed EEG electrode types include wet electrodes, sponge electrodes, or dry electrodes. Wet electrodes are coated with conductive materials, typically Silver/Silver Chloride (Ag/AgCl), utilizing hypertonic electrolyte to build a bridge between the skin and the sensor coating. Sponge electrodes are also usually Ag/AgCl or gold-coated electrodes embedded in sponges saturated with saline water that provides good contact with and serving as the conductive medium between skin and sensor. Finally, dry electrodes require direct contact with the skin beneath the electrode and do not use electrolyte or other fluids to improve conductivity (Lun-De Liao et al., 2012; Niso et al., 2023). Electrodes can further be passive or active (some form of built-in amplification at the electrode level) and their cables can be shielded or non-shielded. However, none of these latter technical specifications seem to lead to differences in data quality when medical-grade wet electrodes are used with appropriate preparation of the recording sites (Scanlon et al., 2021). In general, all sensor types can provide high-quality data in stationary protocols with appropriate preparation of the skin and no active movement of participants. Wet electrodes need longer preparation times and participants’ hair becomes disheveled due to the application of electrolyte or saline water. Additionally, over time, the conductive medium can dry out, resulting in a decline in the quality of the recorded signals. In contrast, dry electrodes generally require less preparation time and involve fewer cleaning demands when compared to wet electrodes. However, dry electrodes tend to exhibit noticeable increases in single-trial and average noise levels, particularly when impedance levels are higher (Chi et al., 2010) and they more often lead to headaches due to the more extensive pressure that is exerted to establish a good skin contact (Mathewson et al., 2017; Zander et al., 2017). Dry electrodes are sometimes also restricted to areas of the head that have less hair to allow a very good and stable connection to the skin and are specifically susceptible to movement of the sensor surface over the skin since no flexible bridge in form of electrolyte is used.

Another important aspect for mobile EEG recordings is the *application and placement of electrodes*. Electrodes are typically incorporated into elastic caps that provide predetermined positions. These positions are standardized across various head sizes, offering a nomenclature for describing activation patterns or effects in a topographic fashion (Oostenveld & Praamstra, 2001). Some commercial systems provide alternative contraptions to hold the electrodes in place, including headbands or flexible arms that can be moved to different sites. The cables connecting the electrode to the amplifier can be loosely arranged or directly incorporated into the cap design prohibiting cable sway. During active behaviors of participants, the way electrodes and cables are fixed is important as specific movements like head turning or walking can lead to movement of the cap, the amplifier, or the electrode cables that then lead to movement-related artifacts in the recorded signal (Gorjan et al., 2022; Gwin et al., 2010; Wunderlich & Gramann, 2021). Rigidity of the electrode holders will lead to movement of the electrodes with increasing acceleration in movements leading to stronger micro-movement of electrodes over the skin that can lead to impedance changes associated with drastic signal distortions. Likewise, the longer and looser the cables are, the more participants’ movement will lead to cable sway which itself can impact the recorded data. In addition, strong cable sway can lead to micro-movement of the electrode itself and lead to the above-described impedance changes introducing noise to the signal. This is especially the case for dry electrodes that are highly susceptible to artifacts during active behaviors of participants.

### EEG Analysis Approaches

#### Data Preprocessing

Beyond the selection of the equipment, the multitude of data analysis methods presents a broad spectrum of analytical possibilities. However, this diversity also introduces a heightened risk of making erroneous decisions, and it’s worth noting that some of these decisions are closely connected to the characteristics of the chosen equipment. In general, due to the volume and capacitive conducted nature of the EEG signal, a varying number of brain and non-brain activities as well as mechanical and electrical artifacts will be recorded at any given moment. This strongly argues for the processing of the recorded data before the extraction of features to avoid interpreting EEG parameters as brain activity when in fact the activity does not or only partially originated from the brain. Especially for mobile EEG recordings that come with increased eye movement, neck, and facial muscle activity as well as potential movement-related mechanical artifacts, an objective and replicable preprocessing of the data is important. Commercial systems often provide only a low number of electrodes and biased distribution of electrode locations together with proprietary algorithms that were validated in stationary protocols and that are unlikely generalizable to mobile settings. When proprietary analysis algorithms are employed, the preprocessing algorithms and the specifications integral to feature extraction cannot undergo critical evaluation, and the results become non-replicable. Hence, results from black-box analysis methods lacking transparency in algorithmic details and the absence of statistics regarding cleaned or interpolated data should be excluded from scientific consideration.

EEG cleaning approaches often focus on the most prominent physiological non-brain contributions to the recorded EEG signal, i.e. eye movement and muscle activity. Usually, several preprocessing steps are applied, including filtering of the signal to remove slow drifts or high-frequency activity that is characteristic of muscle activity. Often, additional time domain cleaning is applied to remove muscle or eye movement-related activity. This can be done in an automated fashion according to predefined criteria or by visual inspection. More specific artifact rejection approaches allow for regressing out eye movement activity (Croft & Barry, 2000) or using spatial filters to filter different non-brain sources from the recorded data without sacrificing samples (Jiang et al., 2019). Regarding data cleaning in mobile EEG recordings, a large number of Neurourbanism studies use Independent Component Analysis (ICA; Makeig et al., 1995), a blind source separation method that allows for the removal of artifactual activity from the recorded signal (Jung et al., 2000). ICA decomposes the data into a matrix of statistically independent source time series with a weight matrix for each resultant independent component (IC) indicating how much each IC contributes to each channel. Using the IC activity time course and spatial distribution allows for differentiating different sources of activity like brain, muscles and eye movement activity. ICs can then be removed from the unmixing matrices and the thus “artifact-cleaned” data can be further processed. While ICA developed into one of the most widely used blind source separation methods for the analyses of EEG data (Delorme & Makeig, 2004), the application of ICA to EEG data requires certain assumptions to be met. Some of the theoretical assumptions include that the extracted processes are stationary, meaning that the statistical properties of the sources and the mixing matrix remain constant over time. Another significant assumption within the ICA model, which bears practical implications for data decomposition, pertains to the necessity of prior knowledge concerning the quantity of underlying sources. In particular, the majority of ICA algorithms partition the data into a quantity of independent components (ICs) equivalent to the number of electrodes, leading to, for instance, 64 ICs in the context of a mobile EEG recording featuring 64 electrodes. Applying ICA to a dataset with 14 electrodes yields only 14 sources to account for the recorded data. If more than 14 sources were active during the recording, the resulting ICs would mix several activation patterns across multiple ICs to elucidate the signal.

Revisiting the physiological underpinnings of the EEG signal, the significance of the number of electrodes becomes apparent for the use of ICA as data cleaning tool. Physiological non-brain sources like eye, facial, and neck muscles contribute significantly through capacitive conduction to the recorded signal. With sufficient number of electrodes, 2 or more ICs will result from the decomposition reflecting vertical and horizontal eye movements while several additional ICs are required to account for the activity of over 40 muscles of the head and neck (Kamibayashi & Richmond, 1998; Westbrook et al., 2023). Beyond potential mechanical artifacts linked to movement and potential electrical artifacts from the environment, the available degrees of freedom to explain the activation of an unknown number of brain processes is markedly diminished. It seems plausible that ICA should not be used with less than 20 channels and researchers need to carefully consider the context and characteristics of their data when applying ICA and interpret the results accordingly since low electrode densities might not allow for separating brain from non-brain activity.

#### EEG Feature extraction

After data cleaning and isolating the time periods of interest in the recorded signal, the relevant features are extracted for further statistical analyses. A feature is commonly defined as a distinctive characteristic or property extracted from biosignals in psychophysiological research, such as the amplitude of an ERP or the spectral power at designated electrode sites. Generally, the EEG signal can be analyzed in the time domain, the frequency domain, and the time-frequency domain (Cohen, 2014; Gramann & Plank, 2019) both on the sensor or the source level after appropriate source reconstruction. Time and time-frequency domain analyses require events, usually derived from controlled stimulus presentations, that allow for extracting stereotypical brain activity associated with the onset of the events, e.g., every time an image is presented on a screen or a response button is pressed. These event-related fluctuations in the EEG signal can then be averaged and analyzed in the time domain as event-related potentials (ERPs; Luck, 2005) or, in the time-frequency domain, as event-related spectral perturbation (ERSPs; Onton & Makeig, 2006). By averaging all time intervals around the onset of events, invariant electrocortical responses to the respective event or class of events are extracted and variable activation in the signal is averaged out. Since event presentation in the real world is difficult to control, the majority of Neurourbanism and Neuro-Architecture studies do not investigate event-related activity. However, innovative analysis approaches allow for investigating event-related activity without stimulus presentation using the behavior of participants itself as event input. For example, eye blinks can be used to extract blink-related potentials from the EEG (Wascher et al., 2014, 2022; Wunderlich & Gramann, 2021) that allow for the comparison of blink-related ERPs in different conditions (e.g., built vs green environment). The electrocardiogram (ECG) can be used to extract R-peaks from the ECG to compute heartbeat-evoked potentials for further investigation (e.g., Luft & Bhattacharya, 2015). In case mobile EEG is combined with eye-tracking, fixation-based ERPs for specific object classes that attract fixations can be investigated also in mobile protocols (e.g., Ladouce et al., 2022).

As an alternative, the transformation of data from the time domain to the frequency domain reveals insights into the energy of specific frequency bands across the entire signal spectrum, contingent upon experimental manipulations. EEG, representing neural oscillatory activity, can be analyzed by decomposing data into weighted sine and cosine functions with different frequencies, phases, and amplitudes. This decomposition is achievable through methods like the Fast Fourier Transform (FFT) converting the signal from the time to the frequency domain. Frequency-domain analyses do not necessitate the controlled presentation of events and can be applied to any time period of interest. Examples include comparing power levels during baseline activity against those in experimental periods or directly contrasting different time intervals contingent on experimental conditions, such as alternating between experiences of urban as compared to green environments in designated experimental blocks. Alterations in power within defined frequency bands, exhibiting a specific topography, have been linked to many psychological constructs, with emotion (for a recent review see Suhaimi et al., 2020) and meditative states (e.g., Cahn & Polich, 2013) attracting strong interest from the Neurourbanism community. The specific topographic distribution of the frequency band of interest can provide important insight into the involved sensory modalities or multimodal integration processes. For example, shifts in visual attention are accompanied by modulations in the posterior alpha band (8–12 Hz) that can be used for monitoring EEG correlates of visual attention (Van Gerven & Jensen, 2009).

Additionally, in moving beyond understanding the brain activity in a single region at the time, the EEG signal can be transformed into the Hilbert space, which provides phase information relevant to timing and synchronization of different brain activities. This allows for instance to understand the coordination of neural activity in different brain regions during different tasks such as perception (Lachaux et al., 1999; Rodriguez et al., 1999). Particularly due to the rhythmic nature of brain function, the very shape of the signals is rich in information (Buzsáki, 2006). For instance, the Hilbert transformed signal can be used to obtain the amplitude envelope of EEG signals. This can be particularly useful for studying amplitude modulation, which reflects the waxing and waning of rhythmic brain activity, in synchronization with the environment (Charalambous & Djebbara, 2023).

A crucial consideration for mobile EEG applications in Neurourbanism research however is the question whether findings regarding spectral power modulations observed in stationary protocols can be extrapolated to mobile recordings without restrictions. This is a particularly intriguing question because numerous patterns of movement-related activity overlap in their frequency bands with frequencies that mirror the cognitive and affective processes under investigation. In addition, the control of potentially confounding factors possible in the lab is mostly absent in the real world, and sensory processing of uncontrolled events will take place in the frequency bands of interest that might be interpreted as power modulation regarding manipulations of the construct under investigation (Barnes et al., 2023).

With this general overview on the requirements of the EEG equipment for recording human brain activity in Neurourbanism, the analyses approaches and extraction of parameters, and their final interpretation, the following section will use these aspects to analyze published papers in the field of Neurourbanism using EEG.

### A Review of Publications in the Field of Neurourbanism Using (Mobile) EEG

A literature review was conducted through the Web of Science (WoS) database and Google Scholar on November 25, 2023, focusing on scientific journal publications exclusively in English. The search in WoS encompassed empirical studies indexed from 1948 onwards, limited to full-length peer-reviewed journal articles. Exclusions were made for reviews, abstracts, and conference proceedings and the main search terms for abstract queries included the following:

AB=((“built environment” OR “architectural space” OR “interior space” OR “environmental design” OR “physical environment” OR “urban” OR "urbanism" OR “urban density” OR "neurourbanism" OR "neuroarchitecture" OR “designed space” OR “urban design” OR “green space” OR “urban landscape”) AND ("EEG" OR “mobile EEG” OR “ambulatory EEG” OR “Mobile Brain/Body Imaging”)).

After the initial search, the results were refined by excluding the search terms “Epilepsy”, “Seizure”, and “Evolutionary Economic Geography”.

The search returned 99 publications of which 55 publications reported empirical EEG research and matched the selection criteria. Based on these results, an additional search was conducted in Google Scholar based on the literature in the identified papers resulting in 75 peer-reviewed papers using EEG in the laboratory or the real world to assess the human brain response to built and natural environments. Nearly 75% of these papers have been published within the last five years. All studies were subsequently classified regarding the mobility score of the EEG system and the participant mobility score according to the classification scheme from Bateson and colleagues (2017). An additional focus was set on the number of electrodes in the selected studies which, together with the system and participant mobility score, allows for a critical evaluation of the reported results regarding potential limitations through movement-related artifacts and restrictions regarding the analysis approach. System scores were derived from the hardware description in the respective studies and participant mobility scores were evaluated based on the description of the study protocol in the selected studies. Since a number of the identified studies used head-mounted virtual reality (VR) allowing restricted or unrestricted movements of the head, it should be noted that the classification from Bateson and colleagues did not further differentiate between walking and head movements with the latter also contributing movement-related artifacts to the recordings (see e.g., Gramann et al., 2010; Jungnickel & Gramann, 2016). For better comparison with previously published classifications the classes were kept as suggested by Bateson (Bateson et al., 2017).

#### Evaluation of selected studies regarding use of Hardware

The studies were evaluated according to the publication guidelines and recommendations for studies using electroencephalography and magnetoencephalography (Keil et al., 2014). Only a subset of the recommended key points was used to allow for a more liberal evaluation of the reviewed studies. All studies were investigated regarding the information on the amplifiers used, including the amplifiers’ type (amplifier name was provided), sampling rate (online sampling rate was stated), and the use of online filters (online filters were described, without having to state the type of filter or roll-off and cut-off parameters). Regarding the specification of electrodes, the criteria included information about the number of electrodes, sensor types (the type of EEG sensor was described, e.g., wet or dry sensor), and sensor locations (sensor locations were specified, including reference electrode(s), providing general location systems like 10% system etc. were considered sufficient). Of note, several studies used consumer market systems that could be researched online regarding these criteria. However, individual configurations might have change standard setups and thus these criteria were considered not fulfilled in case the information was not provided.

The analyses of the selected publications showed that many studies failed to provide sufficient details on amplifier specifications for their recordings, such as the name of the system, the sampling rate and filter settings during recording, and sometimes even the number of electrodes. Nearly all studies (94.7%) reported the amplifier make and 68% of the reviewed studies reported the sample rate during recording. Only 14.7% of all studies reported online filter settings of the amplifier (see Figure 1).

**Figure 1:**
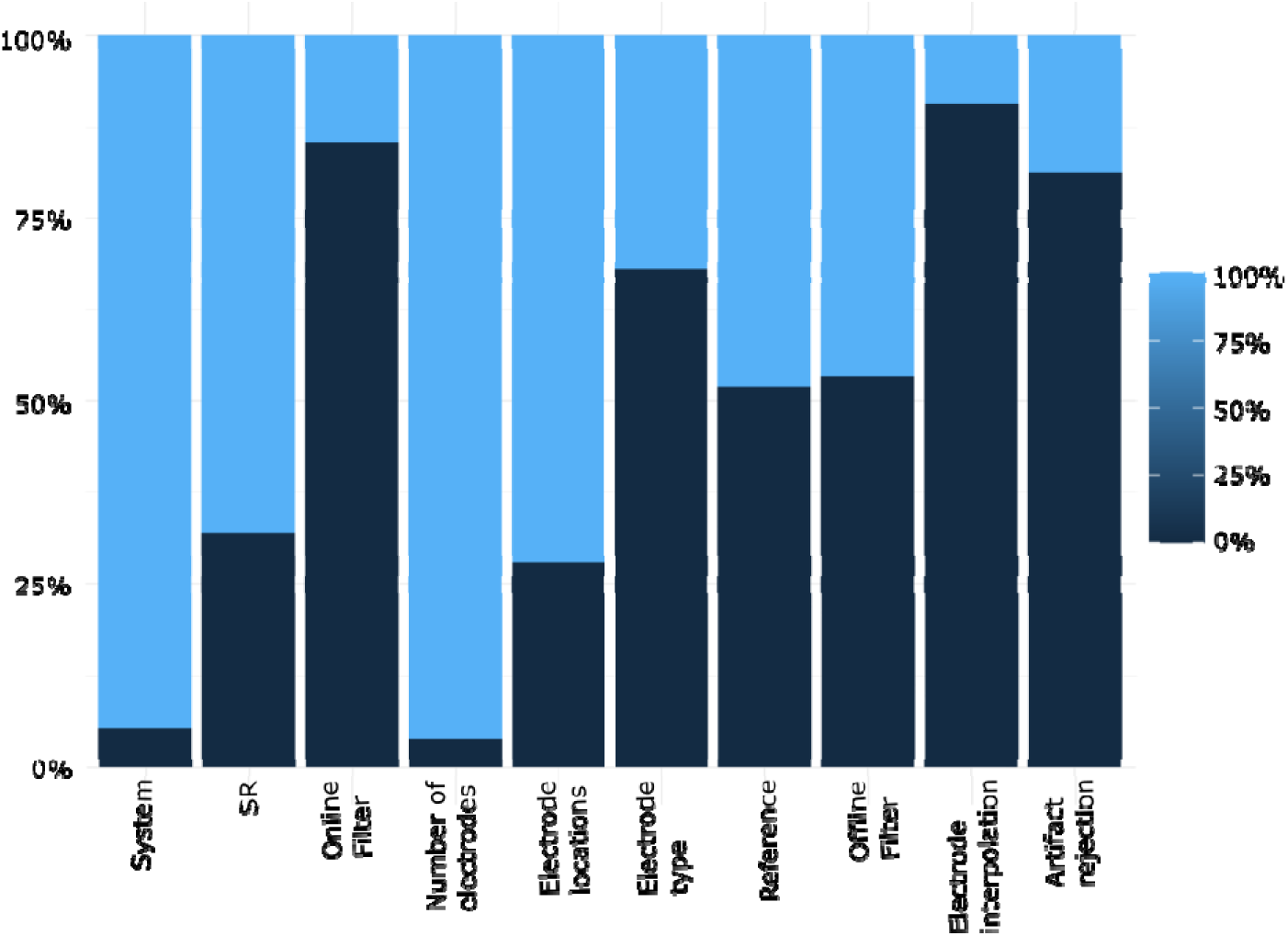
Evaluation criteria applied to reviewed studies. Information on respective category provided in the reviewed studies. Dark filled area displaying the percentage f studies not providing the respective information, light filled are reflecting percentage of studies providing the required information.

Of the 75 studies overall, only two (2.6%) studies failed to report the number of electrodes. The majority of studies (72%) also reported the electrode positions but only 32% of the studies indicated the sensor type (e.g., dry or wet electrodes). Approximately half of the studies (48%) reported which electrode or combination of electrodes was used as the reference. Regarding the data processing criteria (discussed later), the review revealed less than half of the studies (46.7%) to provide information about the filter settings during analyses. Only 18.7% of all studies reported criteria for data rejection and only 9.3% reported whether and how they interpolated artifactual channels.

Overall, a wide range of different systems from diverse companies were used varying in scores for system portability, participant portability as well as the number of electrodes in the respective study (see Figure 2).

**Figure 2:**
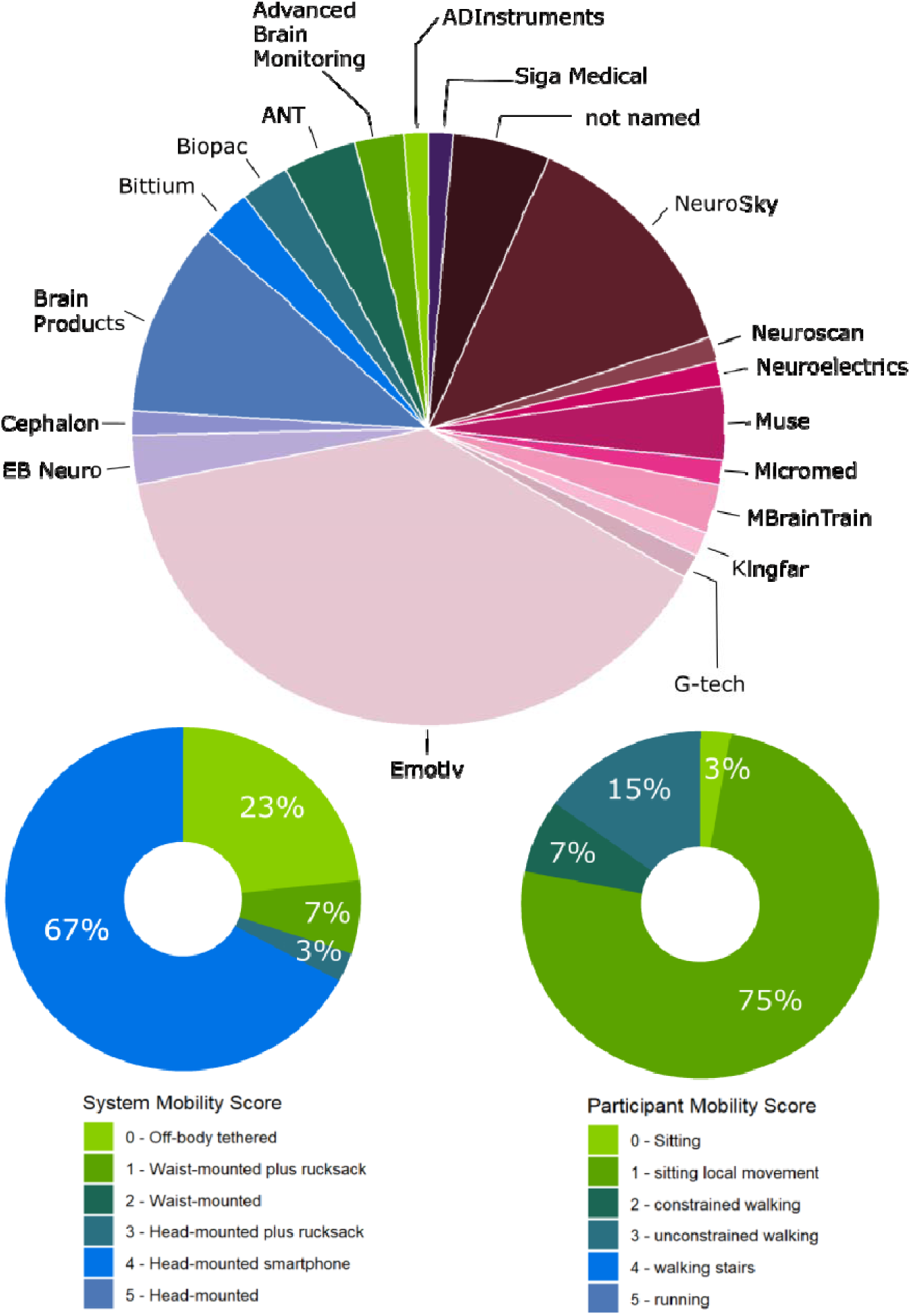
Distribution of EEG systems and setups used in the analyzed Neurourbanism studies categorized according to the CoME score (Bateson et al., 2017). Companies are arranged alphabetically and in counterclockwise direction. Right panel displays distribution of system mobility scores (upper panel) and participant mobility scores (lower panel) across all included studies.

Of the 73 studies that provided information on the number of electrodes, the majority (82.2%) used montages with less than 20 electrodes while 6.8% of the studies used between 22 and 32 electrodes. Further 6 studies used 64 electrodes (8.2%) and 2 studies used 96 or 128 electrodes (each 1.4%). Most of the amplifiers were manufactured by Emotiv (38.7%), NeuroSky (14.7%), and Brain Products (10.7%). In terms of system mobility, as defined by (Bateson et al., 2017), 24% of the amplifier systems had a mobility score of 0, while 7% had a system mobility score of 1, and 3% had a score of 3. With 66%, the majority of EEG systems demonstrated high mobility with a system mobility score of 4. Regarding the participant mobility score, the high percentage of mobile systems used in the studies was not necessarily accompanied by a high participant mobility score in the respective studies. A large number of studies using EEG systems with the amplifiers mounted in the cap still restricted participants’ movements using protocols that required participants to sit still while watching images or movies presented on screens or head-mounted virtual reality displays. Participant mobility scores indicated that 3% of studies involved participants sitting or lying with no movements. The largest portion of all identified studies allowed only minimal movements with participants sitting or lying and responding by button presses or providing other output at the end of experimental trials (76%). Only 7% of the identified studies allowed constrained walking (instructed slow walking or walking on a treadmill), while 14% permitted unconstrained walking (walking at preferred speed in or outside the laboratory).

From Figure 3, it becomes visible that most studies that used EEG systems with a high electrode density used stationary protocols. Only three studies with 64 or more electrodes (with two of the studies reporting different analyses approaches for the same dataset) allowed participants to walk through the (virtual) environment.

**Figure 3:**
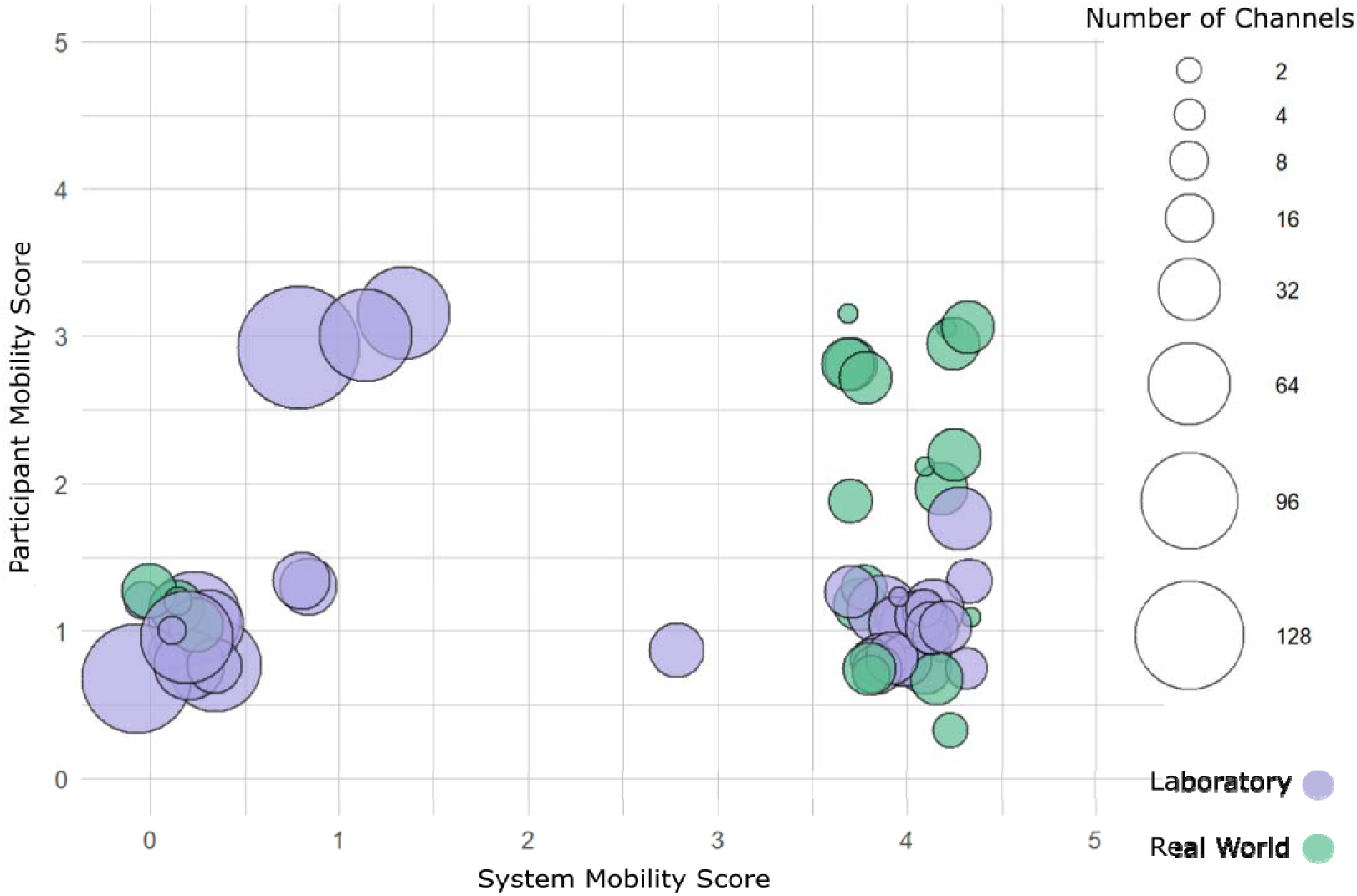
EEG devices used in the identified studies displayed according to their system mobility score an participant mobility score. The number of electrodes is displayed with increasing electrode density indicated by increasing diameter of the respective sphere. Categories indicate electrode number to be smaller or equivalent to the category label (e.g., <= 2 electrodes). Color coding indicates whether the study was conducted in the lab or in the real world. Colors based on colorspace (Zeileis et al., 2019).

**Figure 4:**
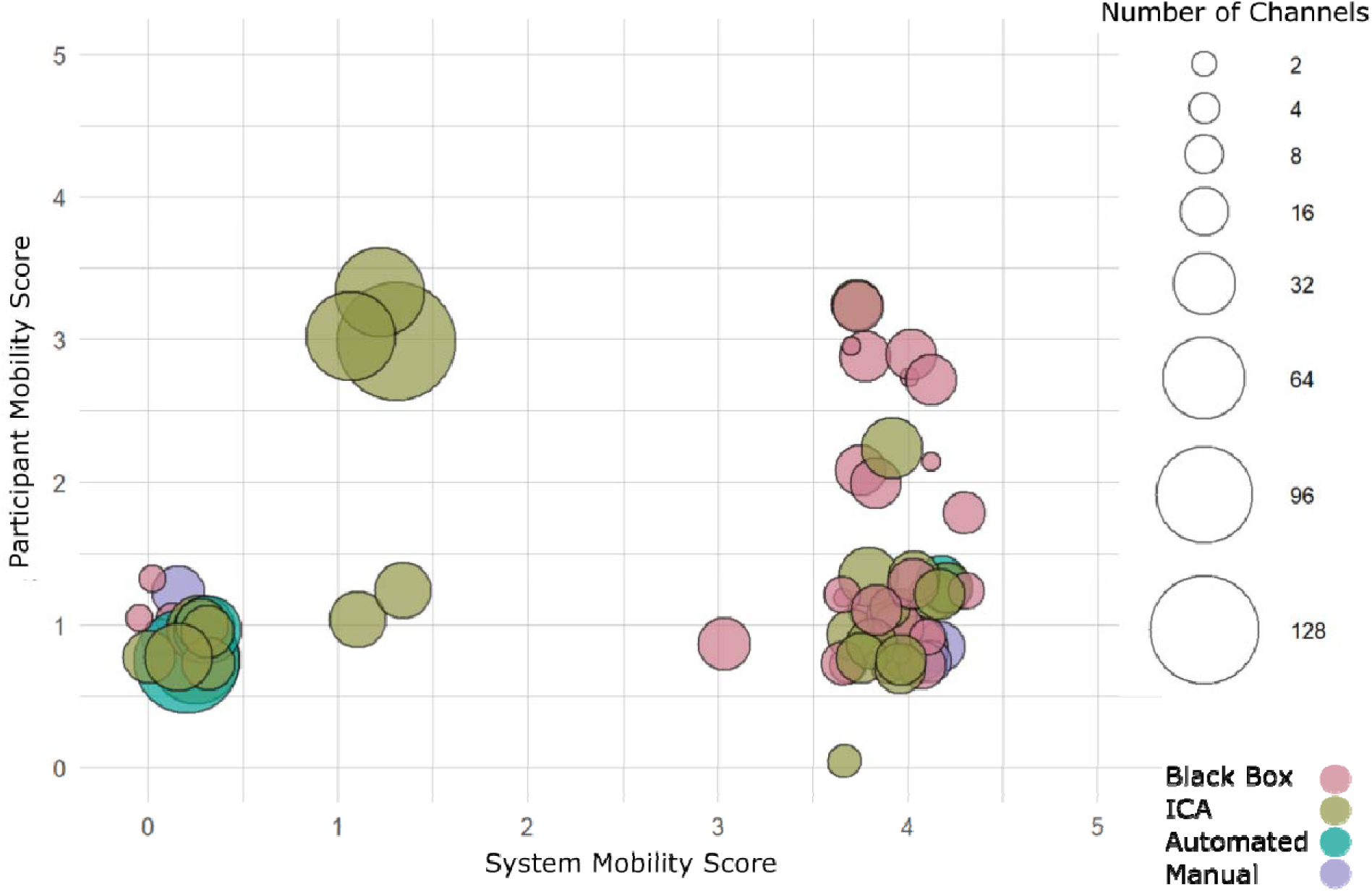
Data analyses approaches for EEG devices used in the identified studies according to their system mobility score and participant mobility score. The number of electrodes is displayed with increasing electrode density indicated by increasing diameter of the respective sphere. Color coding indicates the analyses approach with (pink) Black Box output, (green) ICA, (cyan) automated preprocessing (other than ICA) providing toolbox names and settings, (purple) manual analyses based on visual inspection. Colors based on colorspace (Zeileis et al., 2019).

#### Evaluation of selected studies regarding data preprocessing and Feature Extraction

The following section provides an overview on data preprocessing approaches based on a reduced set of guidelines from the committee report for studies using electroencephalography and magnetoencephalography (Keil et al., 2014). The reduced key point for data processing and feature extraction included information regarding rereferencing (information about the new reference would fulfil the criteria even in case the original reference electrode for recording was not mentioned), the detection of noisy channels and the interpolation of channels (the number of channels that were removed for each participant and the interpolation algorithm used for estimating missing channels were reported) and artifact rejection approaches (artifact rejection procedures are described, including the type and proportion of artifacts rejected). Rereferencing approach were not considered here since the absence of a clear description for rereferencing was only possible in case rereferencing was reported in the first place (see Figure 1). Less than half of the studies (46.7%) provided information about the filter settings and the criteria for data rejection and channel interpolation were provided by only 18.7% and 9.3%, respectively.

Overall, concerning the analysis approaches in the selected studies, only 10 out of 75 studies (13.3%) provided sufficient information to replicate their analysis, while 23 studies (30.7%) offered sufficient details on algorithms and treatment of artifactual data allowing for a partial replication of their results. Alarmingly, 56% of the surveyed studies, i.e. 42 of 75 peer-reviewed and published studies lacked essential details in the methods section or throughout the paper failing to provide information on amplifier make and settings, sample rates during recording or offline down sampling, reference electrodes, filter settings, artifact rejection criteria, or the amount of data removed before feature extraction. This prohibits meaningful assessment of the accuracy of analyses and the validity of the results and interpretation of the data due to insufficient information. These papers are thus unsuitable for consideration in a scientific context. Importantly, among these studies, 33 studies (44%) used proprietary output from commercially available amplifier systems that lack the necessary information to reproduce the results and that do not allow for critical evaluation of the data or analyses approaches and how artifactual data was handled during the preprocessing.

The use of black-box output features was observed mainly for systems with electrode densities below 20 electrodes. This can be attributed mainly to the use of the NeuroSky and Emotiv EEG systems that provide automated classification of EEG data according to different user states.

With 33 studies out of 75, 44% of all identified studies relied on black-box analyses outcome. Beyond the use of non-replicable black box output parameters, 40% (30 studies) of all studies used ICA as an analysis approach to preprocess the data. The use of ICA for some kind of data preprocessing, mostly artifact rejection, is a critical aspect for a large number of the identified studies with less than 20 electrodes due to the limited dimensionality of the data that does not allow dissociating all active brain and non-brain activity patterns. Importantly, the description of the exact use of ICA and subsequent removal of ICs varied drastically. Most of the studies did not provide sufficient information on which ICA algorithm was used, whether the dimensionality was reduced before decomposition, how the resulting ICs were classified as artifactual or functional data or how many ICs were removed, which would be necessary for a replication of the analyses. In one case, ICA was computed on data with 4 electrodes, in a second case ICA was computed on data that was reduced from 14 to 2 electrodes before ICA decomposition without stating how the decomposed data was subsequently used. The largest portion of these studies did not report at all what the outcome of the decomposition was or what was done with the decomposed data. Only 6 studies (8%) used automated or semi-automated data preprocessing before computation of the final features and 6 studies (8%) reported visual inspection of the data for artefact rejection before feature extraction.

With more than 90%, the majority of the studies identified in the present literature search extracted features in the frequency domain. Among those, 8.3% of the studies used both time and frequency domain features and only 2.8% of the studies used only time domain features (ERPs) and 2 studies used source reconstruction or network analyses on the recorded data. The main focus was on alpha power comparing different conditions with 87% of all studies using alpha power alone (alpha or frontal alpha asymmetry) or in combination with other frequency bands.

## Discussion

The present study investigated which EEG systems were used and how the acquired data was analyzed in Neurourbanism research in the laboratory or the real world. A focus regarding aspects of EEG recordings was on the mobility grades regarding the system and the participant mobility as well as the number of electrodes used to acquire the signals. In addition, the processing of the recorded data and the subsequent feature extraction were considered.

The results paint a deeply concerning picture of the scientific rigor in the emerging field of Neurourbanism. More than half of the research papers cannot be replicated and an additional 30% of the residual papers allow only partial replicability, rendering more than 80% of all papers unsuitable for a critical scientific evaluation. Numerous papers lack objectivity in data acquisition, analysis, and result interpretation. Consequently, over 50% of the studies uncovered in this literature review should be categorically excluded from the realm of credible scientific discourse. Alarmingly, these deficient papers were still published despite failing to meet even minimal standards for reporting physiological data collection and analysis. This underscores the imperative need for multidisciplinary expertise in studies of this nature, as relying solely on commercially available systems without scientific competence and critical scrutiny is clearly inadequate. Furthermore, this situation highlights the role of publishers in disseminating misinformation for profit, which seems not exclusive to any single publishing entity. Notably, Elsevier published 26.2% of the studies lacking essential data recording and analysis information, or relying on opaque methodologies that hinder replication. They were followed by MDPI at 19.5%, Taylor and Francis at 11.9%, and Springer at 9.5%. What is particularly alarming is that all of these publishers also offer journals in the field of Neurosciences, which could have facilitated the inclusion of subject matter experts in the editorial process. Regrettably, this does not appear to have occurred.

All studies that were identified in the present work used EEG amplifier technology with good to very good hardware specifications that would allow for recording high quality EEG data. Nonetheless, even though the acquired raw data might have been of very good quality, the raw data was not used in more than half of the studies. The majority used portable amplifier systems reflecting a general trend towards small and lightweight EEG systems even in case these systems were used in stationary protocols that do not necessarily require mobile EEG systems. It seems rather the low-cost aspect than the general mobile use case that these systems provide compared to medical grade or research-grade amplifier systems that render them attractive to researchers confirming previous observations in this direction (Niso et al., 2023). In combination with automated analyses approaches, these systems appear as an attractive alternative for non-experts in the field. Furthermore, exactly these low-cost EEG systems often come with a low number of electrodes with the NeuroSky and Emotive Epoc systems with only 1 or 14 electrodes, respectively, making up the largest portion of systems used in Neurourbanism research. Recording EEG data with only one electrode does not allow for any analytical exclusion of physiological non-brain activity like eye movement or facial muscle contraction and additional movement-related artifacts in case of ambulating participants. This is also still a critical issue for systems with 14 electrodes. Still, none of the reported studies that can be considered non-replicable discussed this problem. Several studies, including some of the studies that were replicable, further claimed that the Emotiv Epoc system was validated for use in the real world by citing a paper from Debener and colleagues (2012), one of the first ambulatory EEG papers in the field. This study, however, did not use the original electrode system of the Emotiv Epoc but replaced the electrodes using medical grade wet electrodes connecting them to the Emotiv amplifier. This is not comparable to the relative inflexible electrode holder system or the quality of the electrodes that come with the off-the-shelf system. As such, the paper by Debener and colleagues did not validate the use of the Emotiv Epoc system in real world settings or protocols with active movements of participants rendering this argument invalid. Moreover, the only study to the knowledge of the author that directly compared a medical grade EEG system with the Emotiv Epoch system in actively moving participants demonstrated significantly decreased data quality for the Emotiv Epoc system (Duvinage et al., 2012). As such, data recorded with any of these low-density systems in mobile protocols should be considered with caution and only appropriate data analyses approaches should be used. ICA is not one of them.

Overall, the electrode density varied significantly across studies. The majority of high-density studies used stationary laboratory protocols and only three of these studies used mobile protocols in the laboratory. Low-density studies used both laboratory as well as real-world protocols with only a minor subset of studies allowing participant movement through the environment. All higher density studies relied on automated data analyses approaches including ICA as a data cleaning tool. In case of high-density recordings, the dimensionality of the acquired data is high enough to account for multiple physiological and mechanical as well as electronic artifacts. In case of low-density recordings, however, this is not the case and the analyses of EEG with 14 or less electrodes might not be adequate to clearly separate brain from non-brain activity. Nonetheless, the majority of studies using low-density recordings used either the output of proprietary black-box algorithms or ICA as a data cleaning tool. What was even more concerning for the studies with low-density recordings that used ICA as preprocessing tool, however, was the description or rather the absence of a description of which ICA algorithm was used, how the data was prepared before ICA decomposition, and how the decomposed data was subsequently used. Only a few studies used toolboxes to automatically label ICs as reflecting artifactual activity, including MARA (Winkler et al., 2011), SASICA (Chaumon et al., 2015), or ICLabel (Pion-Tonachini et al., 2019). A notable observation is that the majority of studies either failed to disclose their approach to IC selection or omitted any discussion about the treatment of decomposed data. This raises concerns regarding the level of transparency and understanding of the analytical methods employed in these studies.

Regarding the *extraction and interpretation of features* from the recorded physiological data, the present review showed that the majority of studies used parameters from the frequency domain with alpha being the most often investigated parameter. Parameter extraction is a crucial step in operationalizing the research question involving the identification and quantification of specific features or characteristics within the data that are relevant to the physiological processes under investigation. However, challenges and problems can arise during the parameter extraction process when the presence of noise and artifacts in the physiological data distort the accuracy of parameter measurements. This is especially problematic, when the experimental protocol leads to non-brain sources contributing to the frequency band of interest. This is the case for eye and head movement-related activity as well as gait-related artifacts in mobile protocols that contribute to a broad frequency range and can distort the signal of interest. Only a small fraction of studies used mobile protocols with the minority providing sufficient electrode density to allow for identification and subsequent removal of eye and body movement-related activity while also missing to report the exact approach that was implemented. None of the low-density studies addressed movement-related activity to potentially impact the extracted parameters at all. The majority of studies identified in the present literature review, however did not use mobile protocols and as such, the extraction and interpretation of frequency parameters might be considered a reliable approach.

Most of the studies in the field of Neurourbanism that recorded EEG data used frequency domain parameters referring to studies from other research domains that describe power modulations in specific frequency bands to covary with the experimental manipulation. Selecting previously established features from different research areas and interpreting these features in the Neurourbanism context is a solid scientific approach. Critically, however, in real-world protocols, there is no control over the onset of events in the environment. Unexpected occurrences, such as individuals abruptly crossing paths or unpredictable variations in traffic noise, environmental sounds like chirping birds or rustling leaves, can introduce uncontrollable factors that impact the moment-to-moment dynamics of human brain activity. As a consequence, the shorter the recording and analyses periods of EEG are, the more likely variation in human brain dynamics will increase due to the impact of uncontrollable events. A direct comparison with features extracted in highly controlled laboratory settings that do not allow for any kind of movement should thus be considered with caution.

The method of stimulus presentation also warrants further scrutiny in this context. Brain activity, inherently responsive to sensory inputs, is particularly relevant in the study of Neurourbanism, a field defined by environmental sensory stimuli. As noted by Eberhard (2009b), the interplay between the brain, behavior, and environment is central to Neurourbanism. This interplay is underscored by the difference in behavioral responses to simulated urban environments (e.g., videos) versus actual urban settings. Considering the profound impact of environmental characteristics on behavior, it is necessary to incorporate complex and contextually rich stimuli in research designs and analyses beyond simplistic representations (e.g., ‘green space’ vs. ‘urban space’) to include multifaceted elements like spatial dimensions, visual boundaries, interactivity, temporal patterns, and other nuanced environmental features.

In summary, a significant portion of EEG studies within the Neurourbanism field exhibited notable shortcomings in scientific rigor and should be excluded from scientific consideration. The principal issues identified in these studies encompassed inadequate reporting of hardware specifications and data acquisition setups, the utilization of a limited number of electrodes for signal recording, often in conjunction with undisclosed proprietary analysis outputs, or inadequate use of Independent Component Analysis (ICA) for data cleaning. When data acquisition and analysis approaches lack objectivity, the generation of reliable and valid results becomes untenable, significantly compromising result interpretation. It is worth noting that individuals lacking expertise in neuroscience occasionally rely on erroneous assumptions concerning EEG, further exacerbated by the inappropriate application of analytical methods to the data that was recorded with questionable quality. All these misconceptions might stem from the available off-the-shelf and black-box EEG systems falsely conveying confidence that “anyone can use EEG”. Should these findings be further extended, generalized, or even suggested as the basis for urban development policies, it becomes essential to implement mechanisms that deter unwarranted extrapolations and their associated consequences. The observation that journals haphazardly publish such research, often prioritizing profit motives over rigorous quality assessments, underscores a significant departure from the established standards of scientific rigor within the scholarly publishing realm, which is expected to serve as a safeguard against such practices.

## Conclusion - Guidelines for (Mobile) EEG Protocols in the Laboratory or the Real World

From the reported problems in the reviewed studies, it seems imperative that the interdisciplinary character of Neurourbanism research becomes indeed interdisciplinary securing expertise from all fields involved in the research question addressed. In case EEG is used to investigate the human brain response to the natural or built environment, an expert from the neuroscience with experience in EEG should be part of the research team as well as experts from the other domains addressed in the specific project.

Specifically with respect to the use of (mobile) EEG in Neurourbanism research, the following guidelines are derived to help researchers in this field to identify the best EEG system and analysis approaches for their study protocols.

### Amplifiers

In general, commercially available amplifiers are adequate for EEG recordings in terms of data quality. However, it’s important to note that many EEG amplifiers on the consumer market are not optimized for protocols involving participants’ free movement. With respect to amplifier specifications, the following points should be considered:

- Most EEG amplifiers that are commercially available provide sufficient bit rates as well

as sample rates allowing for the acquisition of EEG data of good quality. If the frequency domain of interest is limited from 1 to 60 Hz, sampling rates of 128 Hz are sufficient. In case of explorative analyses without upper frequency band limitation, higher sampling rates of 250 Hz or 500 Hz are advised.

- The CMR should be at least 80 dB or higher.
- If mobile recordings are planned, the weight of the amplifier system should be low and the wireless protocol should allow reliable data recording with missed samples being indicated by the system.
- In case no other data streams are synchronized with the EEG, the data can be stored on local memory cards in case the recording does not need any timing information.
- In case additional data streams are recorded together with EEG (e.g., eye-tracking, ECG), synchronization protocols like Lab Streaming Layer (https://github.com/sccn/labstreaminglayer) should be used to allow multimodal data fusion during recording and analysis.

### Number and Distribution of Electrodes

Further challenges arise in (mobile) EEG studies in case of low electrode numbers and uneven distribution of electrodes over the scalp. Insufficient numbers of electrodes and uneven distribution restrict the analyses approaches and feature extraction to the provided recording sites, particularly affecting investigations into potentially yet unknown EEG features. The following guidelines might be considered:

- Never use only one electrode for scientific investigation of human brain activity.
- Always use only one electrode as the reference electrode during recording. The data can always be re-referenced offline to any desired reference setting. The reference electrode should sit tight in the cap and have good contact to the skin (e.g., mastoid references incorporated in the cap are often artifact prone due to the loose fit of the caps next to the ear slits).
- Systems with 14 electrodes might be sufficient in case of stationary protocols when the parameter of interest can be derived from this set of electrodes at the specified electrode positions.
- When participants are allowed to move through the environment, at least 32 but better 64 electrodes or more should be used.
- In case the parameters of interest are known in advance, electrodes should be placed at those locations where the feature is expected based on previous work. In case of explorative studies, the electrodes should be distributed across the entire scalp using standardized electrode locations (e.g., Oostenveld & Praamstra, 2001).
- High numbers of electrodes with even distribution allow for a wider range of analytical approaches (e.g., ICA, source reconstruction, network analyses).

### Kind of Electrodes

In case of stationary protocols without participant movement, most electrode types can provide good-quality data with appropriate preparation of the recording sites. The use of dry electrodes in real-world mobile EEG studies may pose issues with the restriction of electrode locations to sites with less hair often in proximity to the eyes and close to facial muscles. Moreover, dry electrodes are especially artifact-prone during active participant movements due to changes in impedance with movements of the electrode over the skin.

- When mobile protocols are employed, wet electrodes provide less artifact vulnerability and prohibit, to a certain degree, artifacts related to micro-movements of the electrodes.
- With appropriate preparation of the recording sites, passive and active wet electrodes with or without shielding can provide comparable quality (see Scanlon et al., 2021)
- In case of mobile EEG recordings, cables should be fixed in the cap or arranged in a way that cable sway is inhibited as far as possible to avoid movement-related artifacts.

### Data preprocessing

Usually, EEG data is preprocessed to eliminate unwanted, artifactual data. This can be done in a variety of ways each influencing the data in a specific way. The data processing pipelines for stationary data differ from the preprocessing approach for mobile data (Klug & Gramann, 2021). Specific pipelines to preprocess mobile EEG data exist (e.g., Klug et al., 2022). When only time domain features are of interest, filter settings can be used that suppress high and low frequency aspects of the data (e.g., low-pass filter of 40Hz and high-pass filter of 1 Hz) eliminating aspects of muscle activity and slow drifts, respectively. In case frequency domain parameters are of interest, filters that interfere with the frequency bands of interest should be avoided.

- Filter settings depend on the feature of interest; in case of explorative analyses and no specific feature being targeted, filters should be avoided.
- When ICA is used, the model assumptions should be known and met. The quality of the ICA outcome depends on the quality of the recorded data (crap in crap out), the number of potentially active sources, the number of electrodes, and the mobility of participants.
- ICA should not be used with less than 24 electrodes since it is arguably unlikely that the summed number of brain and non-brain sources is lower than 24.
- If lower electrode densities (e.g., 14 electrodes) are used, it is advised to remove only ICs reflecting eye movement as these activity patterns might be well dissociated from other sources due to their high energy in the EEG signal. This, however, does not warrant that no brain activity will be removed too.
- Removing ICs, i,e. eye and muscle activity-related ICs, should have an objective basis that can be reported and replicated (e.g., classification like MARA, ICLabel etc).
- For all recordings but especially for low density recordings, the likelihood of removing functional brain activity increases with increasing number of ICs that are removed from the data.

### Feature extraction and interpretation

Ultimately, the interpretation of EEG data hinges on features observed within specific experimental protocols. These features transition into parameters when their empirical observations are numerically quantified to represent specific aspects of interest, as seen in posterior alpha power to change dependent on the level of relaxation of participants. A psychophysiological indicator, in this context, emerges when a parameter consistently provides reliable information about a targeted phenomenon. In Neurourbanism research, an indicator might be an EEG parameter viewed as a stable representation of an underlying cognitive or emotional state replicated across various protocols and conditions. Nevertheless, such stability for specific parameters is not yet observed in the field.

- Parameters that rely on proprietary algorithms that lack transparency, preventing a thorough evaluation of processing parameters or the reproduction of results should be excluded from scientific investigations.
- Parameters should be interpreted with caution without “cherry-picking” previous studies that support the authors’ claims.
- Parameters that were established in stationary protocols would be interpreted with caution if the experimental protocol included movement.

These guidelines, while not comprehensive, represent key considerations for (mobile) EEG studies drawn from over 15 years of research in mobile brain imaging using EEG. They are intended to assist researchers venturing into this dynamic area of study.

**Table 1:**
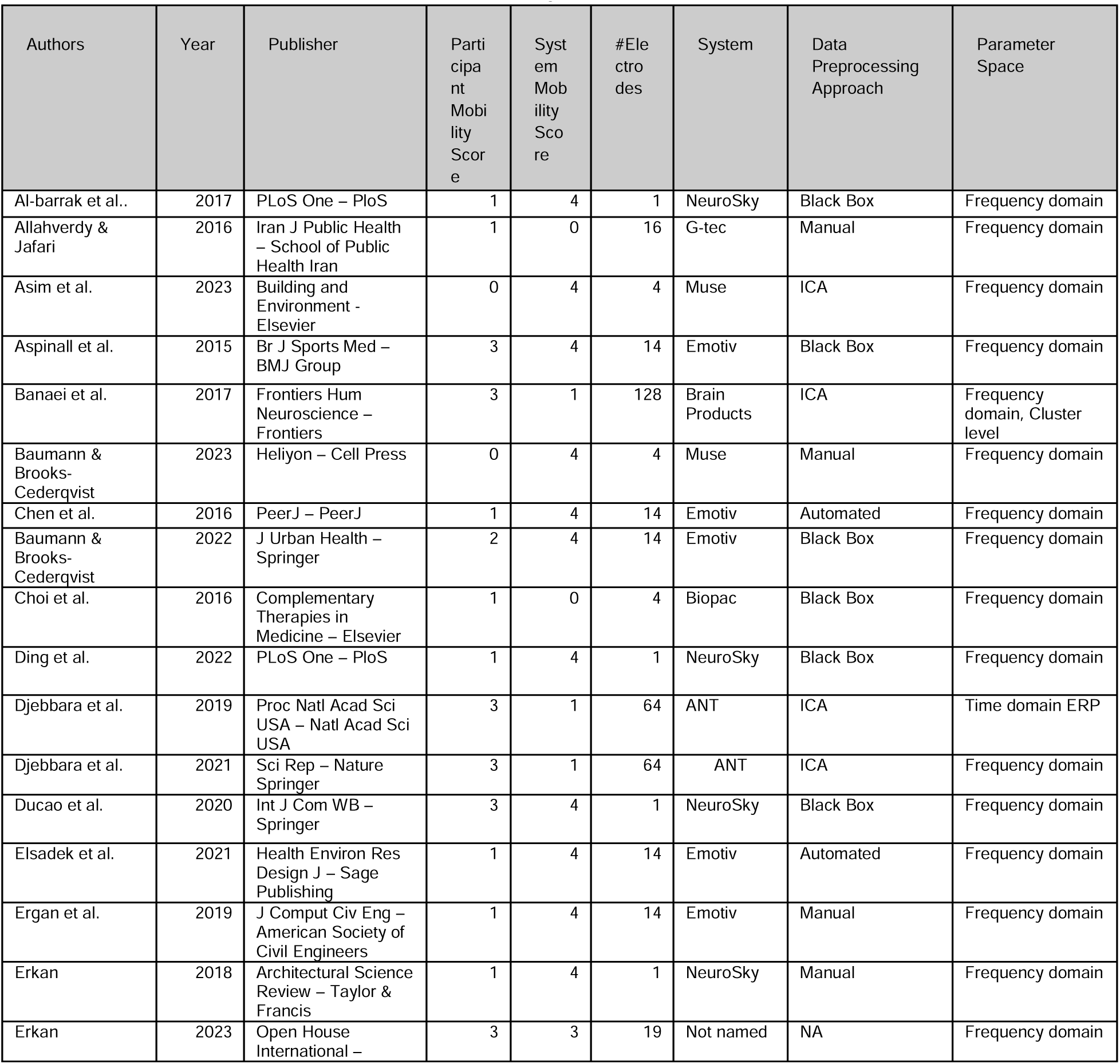
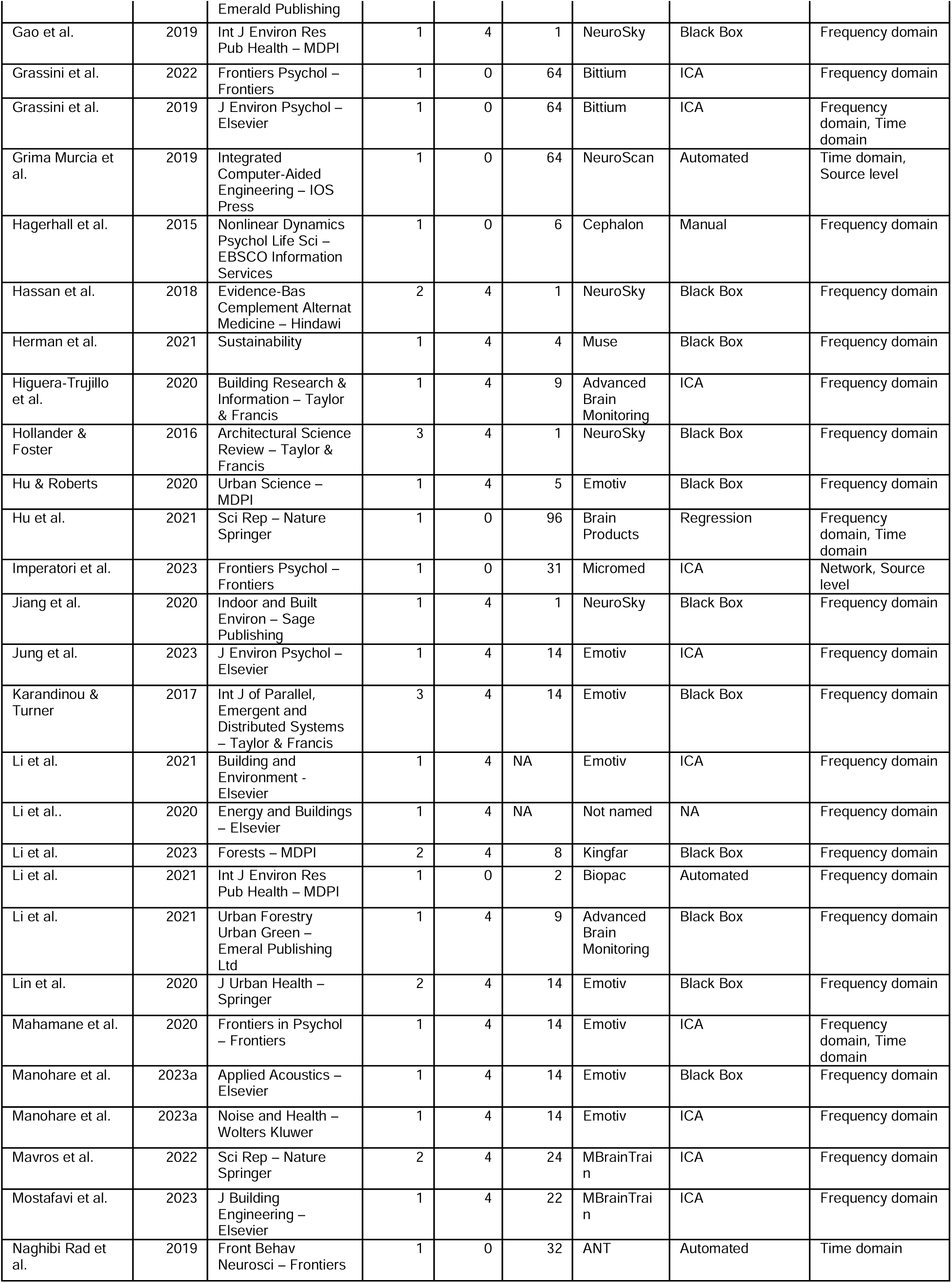
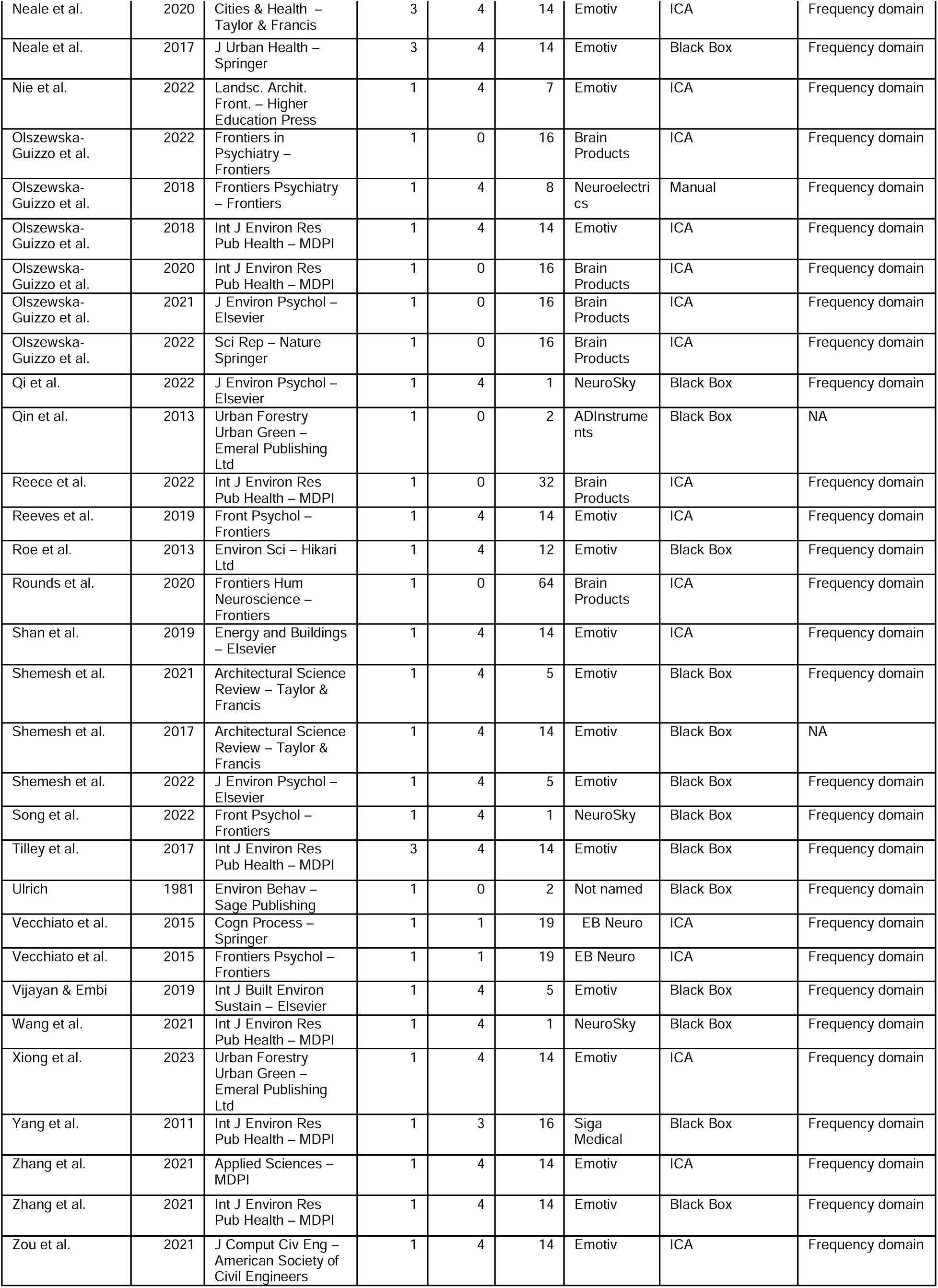
Publications from 1945 to November 2023 using EEG or mobile EEG in Neurourbanism studies.

The author would like to thank Lara Kläffling, Isabelle Sander, Anna Wunderlich, Chris Hilton, and Zak Djebbara for their input and constructive comments preparing this manuscript.

This research was funded under the Excellence Strategy of the Federal Government and the Länder by the Berlin University Alliance.

The author declares he has no conflicts of interest related to this work to disclose.

During the preparation of this work the author used ChatGPT in order to edit sentences and to improve readability of the manuscript. After using this tool/service, the author reviewed and edited the content as needed and takes full responsibility for the content of the publication.

## References

1. Adli, M., Berger, M., Brakemeier, E.-L., Engel, L., Fingerhut, J., & Gomez-Carrillo, A. (2017). Neurourbanism: Towards a new discipline. Lancet Psychiatry, 4, 183–185. 10.1016/S2215-0366(16)30371-6

2. Al-barrak, L., Kanjo, E., & Younis, E. M. G. (2017). NeuroPlace: Categorizing urban places according to mental states. PLOS ONE, 12(9), e0183890. 10.1371/journal.pone.0183890

3. Allahverdy, A., & Jafari, A. H. (2016). Non-auditory Effect of Noise Pollution and Its Risk on Human Brain Activity in Different Audio Frequency Using Electroencephalogram Complexity. Iranian Journal of Public Health, 45(10), 1332–1339.

4. Ancora, L. A., Blanco-Mora, D. A., Alves, I., Bonifácio, A., Morgado, P., & Miranda, B. (2022). Cities and neuroscience research: A systematic literature review. Frontiers in Psychiatry, 13, 983352. 10.3389/fpsyt.2022.983352

5. Asim, F., Chani, P. S., Shree, V., & Rai, S. (2023). Restoring the mind: A neuropsychological investigation of university campus built environment aspects for student well-being. Building and Environment, 244, 110810. 10.1016/j.buildenv.2023.110810

6. Aspinall, P., Mavros, P., Coyne, R., & Roe, J. (2015). The urban brain: Analysing outdoor physical activity with mobile EEG. British Journal of Sports Medicine, 49(4), 272–276. 10.1136/bjsports-2012-091877

7. Banaei, M., Hatami, J., Yazdanfar, A., & Gramann, K. (2017). Walking through Architectural Spaces: The Impact of Interior Forms on Human Brain Dynamics. Frontiers in Human Neuroscience, 11, 477. 10.3389/fnhum.2017.00477

8. Barnes, L., Davidson, M. J., & Alais, D. (2023). *The speed and phase of locomotion dictate saccade probability and simultaneous low-frequency power spectra* [Preprint]. Neuroscience. 10.1101/2023.06.22.546202

9. Bateson, A. D., Baseler, H. A., Paulson, K. S., Ahmed, F., & Asghar, A. U. R. (2017). Categorisation of Mobile EEG: A Researcher’s Perspective. BioMed Research International, 2017, 1–15. 10.1155/2017/5496196

10. Baumann, O., & Brooks-Cederqvist, B. (2023). Multimodal assessment of effects of urban environments on psychological wellbeing. Heliyon, 9(6), e16433. 10.1016/j.heliyon.2023.e16433

11. Buzsáki, G. (2006). Rhythms of the brain. Oxford University Press.

12. Cahn, B. R., & Polich, J. (2013). Meditation states and traits: EEG, ERP, and neuroimaging studies. *Psychology of Consciousness: Theory*, Research, and Practice, 1(S), 48–96. 10.1037/2326-5523.1.S.48

13. Charalambous, E., & Djebbara, Z. (2023). On natural attunement: Shared rhythms between the brain and the environment. Neuroscience & Biobehavioral Reviews, 155, 105438. 10.1016/j.neubiorev.2023.105438

14. Chaumon, M., Bishop, D. V. M., & Busch, N. A. (2015). A practical guide to the selection of independent components of the electroencephalogram for artifact correction. Journal of Neuroscience Methods, 250, 47–63. 10.1016/j.jneumeth.2015.02.025

15. Chen, Z., He, Y., & Yu, Y. (2016). Enhanced functional connectivity properties of human brains during in-situ nature experience. PeerJ, 4, e2210. 10.7717/peerj.2210

16. Chi, Y. M., Jung, T.-P., & Cauwenberghs, G. (2010). Dry-Contact and Noncontact Biopotential Electrodes: Methodological Review. IEEE Reviews in Biomedical Engineering, 3, 106–119. 10.1109/RBME.2010.2084078

17. Choi, J.-Y., Park, S.-A., Jung, S.-J., Lee, J.-Y., Son, K.-C., An, Y.-J., & Lee, S.-W. (2016). Physiological and psychological responses of humans to the index of greenness of an interior space. Complementary Therapies in Medicine, 28, 37–43. 10.1016/j.ctim.2016.08.002

18. Cohen, M. X. (2014). Analyzing Neural Time Series Data: Theory and Practice. MIT Press. https://books.google.de/books?id=rDKkAgAAQBAJ

19. Croft, R. J., & Barry, R. J. (2000). Removal of ocular artifact from the EEG: A review. Neurophysiologie Clinique/Clinical Neurophysiology, 30(1), 5–19. 10.1016/S0987-7053(00)00055-1

20. Debener, S., Minow, F., Emkes, R., Gandras, K., & De Vos, M. (2012). How about taking a low_cost, small, and wireless EEG for a walk? Psychophysiology, 49(11), 1617–1621. 10.1111/j.1469-8986.2012.01471.x

21. Delorme, A., & Makeig, S. (2004). EEGLAB: An open source toolbox for analysis of single-trial EEG dynamics including independent component analysis. Journal of Neuroscience Methods, 134(1), 9–21. 10.1016/j.jneumeth.2003.10.009

22. Dimitrov-Discher, A., Gu, L., Pandit, L., Veer, I. M., Walter, H., Adli, M., & Knöll, M. (2023). Stress and streets: How the network structure of streets is associated with stress-related brain activation. Journal of Environmental Psychology, 91, 102142. 10.1016/j.jenvp.2023.102142

23. Ding, N., Zhong, Y., Li, J., Xiao, Q., Zhang, S., & Xia, H. (2022). Visual preference of plant features in different living environments using eye tracking and EEG. PLOS ONE, 17(12), e0279596. 10.1371/journal.pone.0279596

24. Djebbara, Z., Fich, L. B., & Gramann, K. (2021). The brain dynamics of architectural affordances during transition. Scientific Reports, 11(1), Article 1. 10.1038/s41598-021-82504-w

25. Djebbara, Z., Fich, L. B., Petrini, L., & Gramann, K. (2019). Sensorimotor brain dynamics reflect architectural affordances. Proceedings of the National Academy of Sciences, 116(29), 14769–14778. 10.1073/pnas.1900648116

26. Djebbara, Z., & Kalantari, S. (2023). Affordances and curvature preference: The case of real objects and spaces. Annals of the New York Academy of Sciences, 1527(1), 14–19. 10.1111/nyas.15038

27. Ducao, A., Koen, I., Guo, Z., Frank, J., Willard, C., & Kam, J. (2020). Multimer: Modeling Neurophysiological Experience in Public Urban Space. International Journal of Community Well-Being, 3(4), 465–490. 10.1007/s42413-020-00082-7

28. Duvinage, M., Castermans, T., Dutoit, T., Petieau, M., Hoellinger, T., Saedeleer, C. D., Seetharaman, K., & Cheron, G. (2012). A P300-based Quantitative Comparison between the Emotiv Epoc Headset and a Medical EEG Device. Biomedical Engineering / 765*: Telehealth / 766: Assistive Technologies*. Biomedical Engineering, Innsbruck, Austria. 10.2316/P.2012.764-071

29. Eberhard, J. P. (2009a). Applying Neuroscience to Architecture. Neuron, 62(6), 753–756. 10.1016/j.neuron.2009.06.001

30. Eberhard, J. P. (2009b). Brain landscape: The coexistance of neuroscience and architecture. Oxford University Press.

31. Elsadek, M., Shao, Y., & Liu, B. (2021). Benefits of Indirect Contact With Nature on the Physiopsychological Well-Being of Elderly People. HERD: Health Environments Research & Design Journal, 14(4), 227–241. 10.1177/19375867211006654

32. Ergan, S., Radwan, A., Zou, Z., Tseng, H., & Han, X. (2019). Quantifying Human Experience in Architectural Spaces with Integrated Virtual Reality and Body Sensor Networks. Journal of Computing in Civil Engineering, 33(2), 04018062. 10.1061/(ASCE)CP.1943-5487.0000812

33. Erkan, İ. (2018). Examining wayfinding behaviours in architectural spaces using brain imaging with electroencephalography (EEG). Architectural Science Review, 61(6), 410–428. 10.1080/00038628.2018.1523129

34. Erkan, I. (2023). A neuro-cognitive perspective on urban behavior of people with different moods. Open House International. 10.1108/OHI-10-2022-0252

35. Gao, Zhang, Zhu, Gao, & Qiu. (2019). Exploring Psychophysiological Restoration and Individual Preference in the Different Environments Based on Virtual Reality. International Journal of Environmental Research and Public Health, 16(17), 3102. 10.3390/ijerph16173102

36. Gorjan, D., Gramann, K., De Pauw, K., & Marusic, U. (2022). Removal of movement-induced EEG artifacts: Current state of the art and guidelines. Journal of Neural Engineering, 19(1), 011004. 10.1088/1741-2552/ac542c

37. Gramann, K., Ferris, D. P., Gwin, J., & Makeig, S. (2014). Imaging natural cognition in action. International Journal of Psychophysiology, 91(1), 22–29. 10.1016/j.ijpsycho.2013.09.003

38. Gramann, K., Gwin, J. T., Bigdely-Shamlo, N., Ferris, D. P., & Makeig, S. (2010). Visual Evoked Responses During Standing and Walking. Frontiers in Human Neuroscience, 4. 10.3389/fnhum.2010.00202

39. Gramann, K., Gwin, J. T., Ferris, D. P., Oie, K., Jung, T.-P., Lin, C.-T., Liao, L.-D., & Makeig, S. (2011). Cognition in action: Imaging brain/body dynamics in mobile humans. Reviews in the Neurosciences, 22(6). 10.1515/RNS.2011.047

40. Gramann, K., Jung, T.-P., Ferris, D. P., Lin, C.-T., & Makeig, S. (2014). Toward a new cognitive neuroscience: Modeling natural brain dynamics. Frontiers in Human Neuroscience, 8. 10.3389/fnhum.2014.00444

41. Gramann, K., & Plank, M. (2019). The Use of Electroencephalography in Neuroergonomics. In Neuroergonomics (pp. 11–15). Elsevier. 10.1016/B978-0-12-811926-6.00002-6

42. Grassini, S., Revonsuo, A., Castellotti, S., Petrizzo, I., Benedetti, V., & Koivisto, M. (2019). Processing of natural scenery is associated with lower attentional and cognitive load compared with urban ones. Journal of Environmental Psychology, 62, 1–11. 10.1016/j.jenvp.2019.01.007

43. Grassini, S., Segurini, G. V., & Koivisto, M. (2022). Watching Nature Videos Promotes Physiological Restoration: Evidence From the Modulation of Alpha Waves in Electroencephalography. Frontiers in Psychology, 13, 871143. 10.3389/fpsyg.2022.871143

44. Grima Murcia, M. D., Ortiz, M. J., López-Gordo, M. A., Ferrández, J. M., Sánchez Ferrer, F., & Fernández, E. (2019). Neural representation of different 3D architectural images: An EEG study. Integrated Computer-Aided Engineering, 26(2), 197–205. 10.3233/ICA-180591

45. Gwin, J. T., Gramann, K., Makeig, S., & Ferris, D. P. (2010). Removal of Movement Artifact From High-Density EEG Recorded During Walking and Running. Journal of Neurophysiology, 103(6), 3526–3534. 10.1152/jn.00105.2010

46. Hagerhall, C. M., Laike, T., Küller, M., Marcheschi, E., Boydston, C., & Taylor, R. P. (2015). Human physiological benefits of viewing nature: EEG responses to exact and statistical fractal patterns. Nonlinear Dynamics, Psychology, and Life Sciences, 19(1), 1–12.

47. Hassan, A., Tao, J., Li, G., Jiang, M., Aii, L., Zhihui, J., Zongfang, L., & Qibing, C. (2018). Effects of Walking in Bamboo Forest and City Environments on Brainwave Activity in Young Adults. Evidence-Based Complementary and Alternative Medicine, 2018, e9653857. 10.1155/2018/9653857

48. Herman, K., Ciechanowski, L., & Przegalińska, A. (2021). Emotional Well-Being in Urban Wilderness: Assessing States of Calmness and Alertness in Informal Green Spaces (IGSs) with Muse—Portable EEG Headband. Sustainability, 13(4), 2212. 10.3390/su13042212

49. Higuera-Trujillo, J. L., Llinares Millán, C., Montañana I Aviñó, A., & Rojas, J.-C. (2020). Multisensory stress reduction: A neuro-architecture study of paediatric waiting rooms. Building Research & Information, 48(3), 269–285. 10.1080/09613218.2019.1612228

50. Hollander, J., & Foster, V. (2016). Brain responses to architecture and planning: A preliminary neuro-assessment of the pedestrian experience in Boston, Massachusetts. Architectural Science Review, 59(6), 474–481. 10.1080/00038628.2016.1221499

51. Holleman, G. A., Hooge, I. T. C., Kemner, C., & Hessels, R. S. (2020). The ‘Real-World Approach’ and Its Problems: A Critique of the Term Ecological Validity. Frontiers in Psychology, 11, 721. 10.3389/fpsyg.2020.00721

52. Hu, M., & Roberts, J. (2020). Built Environment Evaluation in Virtual Reality Environments—A Cognitive Neuroscience Approach. Urban Science, 4(4), 48. 10.3390/urbansci4040048

53. Hu, M., Simon, M., Fix, S., Vivino, A. A., & Bernat, E. (2021). Exploring a sustainable building’s impact on occupant mental health and cognitive function in a virtual environment. Scientific Reports, 11(1), 5644. 10.1038/s41598-021-85210-9

54. Imperatori, C., Massullo, C., De Rossi, E., Carbone, G. A., Theodorou, A., Scopelliti, M., Romano, L., Del Gatto, C., Allegrini, G., Carrus, G., & Panno, A. (2023). Exposure to nature is associated with decreased functional connectivity within the distress network: A resting state EEG study. Frontiers in Psychology, 14, 1171215. 10.3389/fpsyg.2023.1171215

55. Jackson, A. F., & Bolger, D. J. (2014). The neurophysiological bases of EEG and EEG measurement: A review for the rest of us. Psychophysiology, 51(11), 1061–1071. 10.1111/psyp.12283

56. Jiang, M., Hassan, A., Chen, Q., & Liu, Y. (2020). Effects of different landscape visual stimuli on psychophysiological responses in Chinese students. Indoor and Built Environment, 29(7), 1006–1016. 10.1177/1420326X19870578

57. Jiang, X., Bian, G.-B., & Tian, Z. (2019). Removal of Artifacts from EEG Signals: A Review. Sensors, 19(5), 987. 10.3390/s19050987

58. Jung, D., Kim, D. I., & Kim, N. (2023). Bringing nature into hospital architecture: Machine learning-based EEG analysis of the biophilia effect in virtual reality. Journal of Environmental Psychology, 89, 102033. 10.1016/j.jenvp.2023.102033

59. Jung, T.-P., Makeig, S., Westerfield, M., Townsend, J., Courchesne, E., & Sejnowski, T. J. (2000). Removal of eye activity artifacts from visual event-related potentials in normal and clinical subjects. Clinical Neurophysiology, 111(10), 1745–1758. 10.1016/S1388-2457(00)00386-2

60. Jungnickel, E., & Gramann, K. (2016). Mobile Brain/Body Imaging (MoBI) of Physical Interaction with Dynamically Moving Objects. Frontiers in Human Neuroscience, 10. 10.3389/fnhum.2016.00306

61. Kamibayashi, L. K., & Richmond, F. J. R. (1998). Morphometry of Human Neck Muscles. Spine, 23(12). https://journals.lww.com/spinejournal/fulltext/1998/06150/morphometry_of_human_neck_muscles.5.aspx

62. Karandinou, A., & Turner, L. (2017). Architecture and neuroscience; what can the EEG recording of brain activity reveal about a walk through everyday spaces? *International Journal of Parallel*, Emergent and Distributed Systems, 32(sup1), S54–S65. 10.1080/17445760.2017.1390089

63. Keil, A., Debener, S., Gratton, G., Junghöfer, M., Kappenman, E. S., Luck, S. J., Luu, P., Miller, G. A., & Yee, C. M. (2014). Committee report: Publication guidelines and recommendations for studies using electroencephalography and magnetoencephalography. Psychophysiology, 51(1), 1–21. 10.1111/psyp.12147

64. Kennedy, D. P., & Adolphs, R. (2011). Stress and the city. Nature, 474(7352), 452–453. 10.1038/474452a

65. Kim, T.-H., Jeong, G.-W., Baek, H.-S., Kim, G.-W., Sundaram, T., Kang, H.-K., Lee, S.-W., Kim, H.-J., & Song, J.-K. (2010). Human brain activation in response to visual stimulation with rural and urban scenery pictures: A functional magnetic resonance imaging study. Science of The Total Environment, 408(12), 2600–2607. 10.1016/j.scitotenv.2010.02.025

66. Klug, M., & Gramann, K. (2021). Identifying key factors for improving ICA_based decomposition of EEG data in mobile and stationary experiments. European Journal of Neuroscience, 54(12), 8406–8420. 10.1111/ejn.14992

67. Klug, M., Jeung, S., Wunderlich, A., Gehrke, L., Protzak, J., Djebbara, Z., Argubi-Wollesen, A., Wollesen, B., & Gramann, K. (2022). *The BeMoBIL Pipeline for automated analyses of multimodal mobile brain and body imaging data* [Preprint]. Neuroscience. 10.1101/2022.09.29.510051

68. Kühn, S., Forlim, C. G., Lender, A., Wirtz, J., & Gallinat, J. (2021). Brain functional connectivity differs when viewing pictures from natural and built environments using fMRI resting state analysis. Scientific Reports, 11(1), 4110. 10.1038/s41598-021-83246-5

69. Lachaux, J.-P., Rodriguez, E., Martinerie, J., & Varela, F. J. (1999). Measuring phase synchrony in brain signals. Human Brain Mapping, 8(4), 194–208. 10.1002/(SICI)1097-0193(1999)8:4<194::AID-HBM4>3.0.CO;2-C

70. Ladouce, S., Mustile, M., Ietswaart, M., & Dehais, F. (2022). Capturing Cognitive Events Embedded in the Real World Using Mobile Electroencephalography and Eye-Tracking. Journal of Cognitive Neuroscience, 34(12), 2237–2255. 10.1162/jocn_a_01903

71. Li, H., Dong, W., Wang, Z., Chen, N., Wu, J., Wang, G., & Jiang, T. (2021). Effect of a Virtual Reality-Based Restorative Environment on the Emotional and Cognitive Recovery of Individuals with Mild-to-Moderate Anxiety and Depression. International Journal of Environmental Research and Public Health, 18(17), 9053. 10.3390/ijerph18179053

72. Li, H., Xie, H., & Woodward, G. (2021). Soundscape components, perceptions, and EEG reactions in typical mountainous urban parks. Urban Forestry & Urban Greening, 64, 127269. 10.1016/j.ufug.2021.127269

73. Li, J., Jin, Y., Lu, S., Wu, W., & Wang, P. (2020). Building environment information and human perceptual feedback collected through a combined virtual reality (VR) and electroencephalogram (EEG) method. Energy and Buildings, 224, 110259. 10.1016/j.enbuild.2020.110259

74. Li, J., Wu, W., Jin, Y., Zhao, R., & Bian, W. (2021). Research on environmental comfort and cognitive performance based on EEG+VR+LEC evaluation method in underground space. Building and Environment, 198, 107886. 10.1016/j.buildenv.2021.107886

75. Li, Y., Zhang, J., Jiang, B., Li, H., & Zhao, B. (2023). Do All Types of Restorative Environments in the Urban Park Provide the Same Level of Benefits for Young Adults? A Field Experiment in Nanjing, China. Forests, 14(7), 1400. 10.3390/f14071400

76. Lin, W., Chen, Q., Jiang, M., Tao, J., Liu, Z., Zhang, X., Wu, L., Xu, S., Kang, Y., & Zeng, Q. (2020). Sitting or Walking? Analyzing the Neural Emotional Indicators of Urban Green Space Behavior with Mobile EEG. Journal of Urban Health, 97(2), 191–203. 10.1007/s11524-019-00407-8

77. Lopes da Silva, F. (2013). EEG and MEG: Relevance to Neuroscience. Neuron, 80(5), 1112– 1128. 10.1016/j.neuron.2013.10.017

78. Luck, S. J. (2005). An introduction to the event-related potential technique. MIT Press.

79. Luft, C. D. B., & Bhattacharya, J. (2015). Aroused with heart: Modulation of heartbeat evoked potential by arousal induction and its oscillatory correlates. Scientific Reports, 5(1), 15717. 10.1038/srep15717

80. Lun-De Liao, Chin-Teng Lin, McDowell, K., Wickenden, A. E., Gramann, K., Tzyy-Ping Jung, Li-Wei Ko, & Jyh-Yeong Chang. (2012). Biosensor Technologies for Augmented Brain– Computer Interfaces in the Next Decades. Proceedings of the IEEE, 100(Special Centennial Issue), 1553–1566. 10.1109/JPROC.2012.2184829

81. Mahamane, S., Wan, N., Porter, A., Hancock, A. S., Campbell, J., Lyon, T. E., & Jordan, K. E. (2020). Natural Categorization: Electrophysiological Responses to Viewing Natural Versus Built Environments. Frontiers in Psychology, 11. https://www.frontiersin.org/articles/10.3389/fpsyg.2020.00990

82. Makeig, S., Bell, A. J., Jung, T.-P., & Sejnowski, T. J. (1995). Independent component analysis of electroencephalographic data. Advances in Neural Information Processing Systems, 8.

83. Makeig, S., Gramann, K., Jung, T.-P., Sejnowski, T. J., & Poizner, H. (2009). Linking brain, mind and behavior. International Journal of Psychophysiology, 73(2), 95–100. 10.1016/j.ijpsycho.2008.11.008

84. Manohare, M., Rajasekar, E., & Parida, M. (2023a). Analysing the Change in Brain Waves due to Heterogeneous Road Traffic Noise Exposure Using Electroencephalography Measurements. Noise and Health, 25(116), 36–54.

85. Manohare, M., Rajasekar, E., & Parida, M. (2023b). Electroencephalography based classification of emotions associated with road traffic noise using Gradient boosting algorithm. Applied Acoustics, 206, 109306. 10.1016/j.apacoust.2023.109306

86. Mathewson, K. E., Harrison, T. J. L., & Kizuk, S. A. D. (2017). High and dry? Comparing active dry EEG electrodes to active and passive wet electrodes. Psychophysiology, 54(1), 74–82. 10.1111/psyp.12536

87. Mavros, P., J Wälti, M., Nazemi, M., Ong, C. H., & Hölscher, C. (2022). A mobile EEG study on the psychophysiological effects of walking and crowding in indoor and outdoor urban environments. Scientific Reports, 12(1), Article 1. 10.1038/s41598-022-20649-y

88. Mehta, R. K., & Parasuraman, R. (2013). Neuroergonomics: A review of applications to physical and cognitive work. Frontiers in Human Neuroscience, 7. 10.3389/fnhum.2013.00889

89. Mostafavi, A., Cruz-Garza, J. G., & Kalantari, S. (2023). Enhancing lighting design through the investigation of illuminance and correlated color Temperature’s effects on brain activity: An EEG-VR approach. Journal of Building Engineering, 75, 106776. 10.1016/j.jobe.2023.106776

90. Naghibi Rad, P., Shahroudi, A. A., Shabani, H., Ajami, S., & Lashgari, R. (2019). Encoding Pleasant and Unpleasant Expression of the Architectural Window Shapes: An ERP Study. Frontiers in Behavioral Neuroscience, 13, 186. 10.3389/fnbeh.2019.00186

91. Neale, C., Aspinall, P., Roe, J., Tilley, S., Mavros, P., Cinderby, S., Coyne, R., Thin, N., Bennett, G., & Thompson, C. W. (2017). The Aging Urban Brain: Analyzing Outdoor Physical Activity Using the Emotiv Affectiv Suite in Older People. Journal of Urban Health, 94(6), 869–880. 10.1007/s11524-017-0191-9

92. Neale, C., Aspinall, P., Roe, J., Tilley, S., Mavros, P., Cinderby, S., Coyne, R., Thin, N., & Ward Thompson, C. (2020). The impact of walking in different urban environments on brain activity in older people. Cities & Health, 4(1), 94–106. 10.1080/23748834.2019.1619893

93. Nie, W., Jia, J., Wang, M., Sun, J., & Li, G. (2022). Research on the Impact of Panoramic Green View Index of Virtual Reality Environments on Individuals’ Pleasure Level Based on EEG Experiment. Landscape Architecture Frontiers, 10(2), 36. 10.15302/J-LAF-1-020059

94. Niso, G., Romero, E., Moreau, J. T., Araujo, A., & Krol, L. R. (2023). Wireless EEG: A survey of systems and studies. NeuroImage, 269, 119774. 10.1016/j.neuroimage.2022.119774

95. Nunez, P. L., & Srinivasan, R. (2006). Electric fields of the brain: The neurophysics of EEG (2nd ed). Oxford University Press.

96. Olszewska-Guizzo, A. A., Paiva, T. O., & Barbosa, F. (2018). Effects of 3D Contemplative Landscape Videos on Brain Activity in a Passive Exposure EEG Experiment. Frontiers in Psychiatry, 9. https://www.frontiersin.org/articles/10.3389/fpsyt.2018.00317

97. Olszewska-Guizzo, A., Escoffier, N., Chan, J., & Puay Yok, T. (2018). Window View and the Brain: Effects of Floor Level and Green Cover on the Alpha and Beta Rhythms in a Passive Exposure EEG Experiment. International Journal of Environmental Research and Public Health, 15(11), 2358. 10.3390/ijerph15112358

98. Olszewska-Guizzo, A., Fogel, A., Escoffier, N., Sia, A., Nakazawa, K., Kumagai, A., Dan, I., & Ho, R. (2022). Therapeutic Garden With Contemplative Features Induces Desirable Changes in Mood and Brain Activity in Depressed Adults. Frontiers in Psychiatry, 13, 757056. 10.3389/fpsyt.2022.757056

99. Olszewska-Guizzo, A., Sia, A., Fogel, A., & Ho, R. (2020). Can Exposure to Certain Urban Green Spaces Trigger Frontal Alpha Asymmetry in the Brain?—Preliminary Findings from a Passive Task EEG Study. International Journal of Environmental Research and Public Health, 17(2), 394. 10.3390/ijerph17020394

100. Onton, J., & Makeig, S. (2006). Information-based modeling of event-related brain dynamics. In Progress in Brain Research (Vol. 159, pp. 99–120). Elsevier. 10.1016/S0079-6123(06)59007-7

101. Oostenveld, R., & Praamstra, P. (2001). The five percent electrode system for high-resolution EEG and ERP measurements. Clinical Neurophysiology, 112(4), 713–719. 10.1016/S1388-2457(00)00527-7

102. Pion-Tonachini, L., Kreutz-Delgado, K., & Makeig, S. (2019). The ICLabel dataset of electroencephalographic (EEG) independent component (IC) features. Data in Brief, 25, 104101. 10.1016/j.dib.2019.104101

103. Qi, Y., Chen, Q., Lin, F., Liu, Q., Zhang, X., Guo, J., Qiu, L., & Gao, T. (2022). Comparative study on birdsong and its multi-sensory combinational effects on physio-psychological restoration. Journal of Environmental Psychology, 83, 101879. 10.1016/j.jenvp.2022.101879

104. Qin, J., Zhou, X., Sun, C., Leng, H., & Lian, Z. (2013). Influence of green spaces on environmental satisfaction and physiological status of urban residents. Urban Forestry & Urban Greening, 12(4), 490–497. 10.1016/j.ufug.2013.05.005

105. Reece, R., Bornioli, A., Bray, I., & Alford, C. (2022). Exposure to Green and Historic Urban Environments and Mental Well-Being: Results from EEG and Psychometric Outcome Measures. International Journal of Environmental Research and Public Health, 19(20), Article 20. 10.3390/ijerph192013052

106. Reeves, J. P., Knight, A. T., Strong, E. A., Heng, V., Neale, C., Cromie, R., & Vercammen, A. (2019). The Application of Wearable Technology to Quantify Health and Wellbeing Co-benefits From Urban Wetlands. Frontiers in Psychology, 10, 1840. 10.3389/fpsyg.2019.01840

107. Rodriguez, E., George, N., Lachaux, J.-P., Martinerie, J., Renault, B., & Varela, F. J. (1999). Perception’s shadow: Long-distance synchronization of human brain activity. Nature, 397(6718), 430–433. 10.1038/17120

108. Roe, J. J., Aspinall, P. A., Mavros, P., & Coyne, R. (2013). Engaging the brain: The impact of natural versus urban scenes using novel EEG methods in an experimental setting. Environmental Sciences, 1, 93–104. 10.12988/es.2013.3109

109. Rounds, J. D., Cruz-Garza, J. G., & Kalantari, S. (2020). Using Posterior EEG Theta Band to Assess the Effects of Architectural Designs on Landmark Recognition in an Urban Setting. Frontiers in Human Neuroscience, 14. https://www.frontiersin.org/articles/10.3389/fnhum.2020.584385

110. Scanlon, J. E. M., Jacobsen, N. S. J., Maack, M. C., & Debener, S. (2021). Does the electrode amplification style matter? A comparison of active and passive EEG system configurations during standing and walking. European Journal of Neuroscience, 54(12), 8381–8395. 10.1111/ejn.15037

111. Shamay-Tsoory, S. G., & Mendelsohn, A. (2019). Real-Life Neuroscience: An Ecological Approach to Brain and Behavior Research. Perspectives on Psychological Science, 14(5), 841–859. 10.1177/1745691619856350

112. Shan, X., Yang, E.-H., Zhou, J., & Chang, V. W. C. (2019). Neural-signal electroencephalogram (EEG) methods to improve human-building interaction under different indoor air quality. Energy and Buildings, 197, 188–195. 10.1016/j.enbuild.2019.05.055

113. Shemesh, A., Leisman, G., Bar, M., & Grobman, Y. J. (2021). A neurocognitive study of the emotional impact of geometrical criteria of architectural space. Architectural Science Review, 64(4), 394–407. 10.1080/00038628.2021.1940827

114. Shemesh, A., Leisman, G., Bar, M., & Grobman, Y. J. (2022). The emotional influence of different geometries in virtual spaces: A neurocognitive examination. Journal of Environmental Psychology, 81, 101802. 10.1016/j.jenvp.2022.101802

115. Shemesh, A., Talmon, R., Karp, O., Amir, I., Bar, M., & Grobman, Y. J. (2017). Affective response to architecture – investigating human reaction to spaces with different geometry. Architectural Science Review, 60(2), 116–125. 10.1080/00038628.2016.1266597

116. Song, R., Chen, Q., Zhang, Y., Jia, Q., He, H., Gao, T., & Qiu, L. (2022). Psychophysiological restorative potential in cancer patients by virtual reality (VR)-based perception of natural environment. Frontiers in Psychology, 13, 1003497. 10.3389/fpsyg.2022.1003497

117. Spiers, H. J., & Maguire, E. A. (2007). Decoding human brain activity during real-world experiences. Trends in Cognitive Sciences, 11(8), 356–365. 10.1016/j.tics.2007.06.002

118. Suhaimi, N. S., Mountstephens, J., & Teo, J. (2020). EEG-Based Emotion Recognition: A State-of-the-Art Review of Current Trends and Opportunities. Computational Intelligence and Neuroscience, 2020, 1–19. 10.1155/2020/8875426

119. Tilley, S., Neale, C., Patuano, A., & Cinderby, S. (2017). Older People’s Experiences of Mobility and Mood in an Urban Environment: A Mixed Methods Approach Using Electroencephalography (EEG) and Interviews. International Journal of Environmental Research and Public Health, 14(2), Article 2. 10.3390/ijerph14020151

120. Ulrich, R. S. (1981). Natural Versus Urban Scenes: Some Psychophysiological Effects. Environment and Behavior, 13(5), 523–556. 10.1177/0013916581135001

121. United Nations Department of Economic and Social Affairs. (2019). World Urbanization Prospects: The 2018 Revision. UN. 10.18356/b9e995fe-en

122. Vallet, W., & Van Wassenhove, V. (2023). Can cognitive neuroscience solve the lab-dilemma by going wild? Neuroscience & Biobehavioral Reviews, 155, 105463. 10.1016/j.neubiorev.2023.105463

123. Van Gerven, M., & Jensen, O. (2009). Attention modulations of posterior alpha as a control signal for two-dimensional brain–computer interfaces. Journal of Neuroscience Methods, 179(1), 78–84. 10.1016/j.jneumeth.2009.01.016

124. Vecchiato, G., Jelic, A., Tieri, G., Maglione, A. G., De Matteis, F., & Babiloni, F. (2015). Neurophysiological correlates of embodiment and motivational factors during the perception of virtual architectural environments. Cognitive Processing, 16(S1), 425–429. 10.1007/s10339-015-0725-6

125. Vecchiato, G., Tieri, G., Jelic, A., De Matteis, F., Maglione, A. G., & Babiloni, F. (2015). Electroencephalographic Correlates of Sensorimotor Integration and Embodiment during the Appreciation of Virtual Architectural Environments. Frontiers in Psychology, 6. 10.3389/fpsyg.2015.01944

126. Vijayan, V. T., & Embi, M. R. (2019). Probing Phenomenological Experiences Through Electroencephalography Brainwave Signals In Neuroarchitecture Study. International Journal of Built Environment and Sustainability, 6(3), 11–20. 10.11113/ijbes.v6.n3.360

127. Wang, Y., Qu, H., Bai, T., Chen, Q., Li, X., Luo, Z., Lv, B., & Jiang, M. (2021). Effects of Variations in Color and Organ of Color Expression in Urban Ornamental Bamboo Landscapes on the Physiological and Psychological Responses of College Students. International Journal of Environmental Research and Public Health, 18(3), 1151. 10.3390/ijerph18031151

128. Wascher, E., Arnau, S., Gutberlet, M., Chuang, L. L., Rinkenauer, G., & Reiser, J. E. (2022). Visual Demands of Walking Are Reflected in Eye-Blink-Evoked EEG-Activity. Applied Sciences, 12(13), 6614. 10.3390/app12136614

129. Wascher, E., Heppner, H., & Hoffmann, S. (2014). Towards the measurement of event-related EEG activity in real-life working environments. International Journal of Psychophysiology, 91(1), 3–9. 10.1016/j.ijpsycho.2013.10.006

130. Westbrook, K. E., Nessel, T. A., Hohman, M. H., & Varacallo, M. (2023). Anatomy, Head and Neck: Facial Muscles. In StatPearls. StatPearls Publishing. http://www.ncbi.nlm.nih.gov/books/NBK493209/

131. Winkler, I., Haufe, S., & Tangermann, M. (2011). Automatic Classification of Artifactual ICA-Components for Artifact Removal in EEG Signals. Behavioral and Brain Functions, 7(1), 30. 10.1186/1744-9081-7-30

132. Wirth, L. (1938). Urbanism as a Way of Life. American Journal of Sociology, 44(1), 1–24. 10.1086/217913

133. Wunderlich, A., & Gramann, K. (2021). Eye movement_related brain potentials during assisted navigation in real_world environments. European Journal of Neuroscience, 54(12), 8336–8354. 10.1111/ejn.15095

134. Xiong, X., Jin, H., Hu, W., Zeng, C., Huang, Q., Cui, X., Zhang, M., & Jin, Y. (2023). Benefits of Jasminum polyanthum’s natural aromas on human emotions and moods. Urban Forestry & Urban Greening, 86, 128010. 10.1016/j.ufug.2023.128010

135. Yang, F., Bao, Z. Y., & Zhu, Z. J. (2011). An Assessment of Psychological Noise Reduction by Landscape Plants. International Journal of Environmental Research and Public Health, 8(4), 1032–1048. 10.3390/ijerph8041032

136. Zander, T. O., Andreessen, L. M., Berg, A., Bleuel, M., Pawlitzki, J., Zawallich, L., Krol, L. R., & Gramann, K. (2017). Evaluation of a Dry EEG System for Application of Passive Brain-Computer Interfaces in Autonomous Driving. Frontiers in Human Neuroscience, 11. 10.3389/fnhum.2017.00078

137. Zeileis, A., Fisher, J. C., Hornik, K., Ihaka, R., McWhite, C. D., Murrell, P., Stauffer, R., & Wilke, C. O. (2019). *colorspace: A Toolbox for Manipulating and Assessing Colors and Palettes* (arXiv:1903.06490). arXiv. http://arxiv.org/abs/1903.06490

138. Zhang, S., Feng, X., & Shen, Y. (2021). Quantifying Auditory Presence Using Electroencephalography. Applied Sciences, 11(21), 10461. 10.3390/app112110461

139. Zhang, Z., Zhuo, K., Wei, W., Li, F., Yin, J., & Xu, L. (2021). Emotional Responses to the Visual Patterns of Urban Streets: Evidence from Physiological and Subjective Indicators. International Journal of Environmental Research and Public Health, 18(18), 9677. 10.3390/ijerph18189677

140. Zou, Z., Ergan, S., Fisher-Gewirtzman, D., & Curtis, C. (2021). Quantifying the Impact of Urban Form on Human Experience: Experiment Using Virtual Environments and Electroencephalogram. Journal of Computing in Civil Engineering, 35(3), 04021004. 10.1061/(ASCE)CP.1943-5487.0000966

